# Development of sensorimotor responses in larval zebrafish: a comparison between wild-type and GCaMP6s transgenic line

**DOI:** 10.1101/2024.09.10.612316

**Authors:** Margaux Caperaa, Mathilde Roland-Caverivière, Chelsea Herdman, Nesrine Imloul, Sandrine Poulin, Mado Lemieux, Paul De Koninck, Gabriel D Bossé

**Author notes:** **Correspondence:** Gabriel Bossé.

## Abstract

During early development, zebrafish larvae exhibit stereotypical behaviors, which rapidly become more complex. Thus, the generation of mutant transgenic lines that maintain transparency throughout their larval stage and that can be used to record brain activity has offered strategic opportunities to investigate the underlying neural correlates of behavior establishment. However, few studies have documented the behavioral profile of these lines during larval development. Here, we set up a behavioral characterization using diverse stimuli (light and vibration) throughout larval development to compare the responses of a transgenic strain expressing a pan-neuronal calcium indicator (GCaMP6s) with that of a wild-type strain. Interestingly, we report a drastic switch in behavioral responses to light transitions at 11 days post-fertilization (dpf) and to vibration stimuli at 14 dpf in both lines. These data highlight a specific time window of behavioral complexification. Meanwhile, we found no major difference in the maturation of sensorimotor responses between GCaMP6s and wild-type strains. Thus, these results support using GCaMP6s strain in investigating the neural mechanisms underlying the developmental maturation of sensorimotor responses. We observed nevertheless some minor differences that suggest careful attention should be taken when using mutant/transgenic lines for behavioral studies.

**Highlights:**

- Longitudinal investigation of sensorimotor responses by zebrafish during their larval development
- During the second week of development, larval zebrafish switch their motor response to light transition
- Pan-neuronal nuclear expression of GCaMP6s has little impact on larval fish response to various stimuli

## 1. Introduction

Zebrafish have become an increasingly popular model in the field of behavioral neuroscience. They have been used to study a range of behaviors, from simple optomotor responses to more complex behaviors, such as reward-based conditioning and social interactions [1–4]. As a result of the increased adoption of this model, new and innovative methods have been developed to investigate the neuronal networks that underlie these behaviors.

At the larval stage, zebrafish exhibit particularly attractive characteristics for neuroscience research. i) They express diverse behaviors for instance, as early as 3-5 days post-fertilization (dpf), zebrafish larvae display various sensorimotor responses, non-associative memory (habituation), and fear responses[5]. Their behavioral repertoire is further enriched over the first few weeks of development, establishing more complex behaviors, such as associative memory and social interactions[6–8]. ii) For drug discovery, they develop a functional blood-brain barrier by 3-5 dpf[9]. At that age, they are small enough to test in a 96-well plate format, enabling large-scale small molecule screening and disease[10–12]. iii) In recent years, the development of nearly transparent fish lines by mutating pigment genes, such as the *casper* line, has enabled the development of advanced optical imaging approaches without the need to suppress pigment formation [13]. Despite the rise in popularity of zebrafish, longitudinal studies investigating behavior across development remain scarce. Moreover, the behavioral responses of older larvae and juvenile zebrafish have been less well characterized.

One of the most significant breakthroughs in zebrafish neurobiology was the development of transgenic fish expressing a genetically encoded calcium (Ca^2+^) indicator, GCaMP, in the *casper* mutant. This line tg(Elavl3:H2B-GCaMP6s) expresses GCaMP6s in the nucleus of all neurons, enabling unprecedented brain-wide functional imaging, with single-cell resolution, in a vertebrate model[14, 15]. It has been utilized to study neuronal visuomotor responses, disease modelling and memory formation[16–18]. Recent improvements in live microscopy protocols have shown that functional imaging is not restricted to early larvae stages. It can be performed in larvae as mature as 3-4 weeks old[19, 20], paving the way for investigating neuronal networks responsible for more intricate behaviors, such as associative memory.

As zebrafish neurobiology gains momentum, behavioral assays will inevitably require the use of genetically modified lines, raising the question of whether differences in behavioral outcomes might arise in comparison to wild-type fish. One notable study by *Parker et al*.[21] found that the *casper* strain did not exhibit any difference in anxiety, shoaling and preference for nicotine at the adult stage compared to a control line (*Tu* line). Additionally, there were no noticeable differences in baseline anxiety levels between wild-type (WT), *casper*, and transgenic fish expressing a tissue-specific eGFP at the adult stage[22].

Few studies have examined whether the expression of GCaMP in neurons throughout the body affects the behavior of larvae. A study by Yang et al.[23] reported that Elavl3:H2B-GCaMP6f transgenic zebrafish larvae (7-10 dpf) demonstrated the same abilities as WT-AB fish in an operant learning task.

Motor activity and photomotor responses in zebrafish larvae are well established in wild-type lines[24, 25]. To identify potential alterations in transgenic tg(Elavl3:H2B-GCaMP6s) fish, we compared its behavioral responses to wild-type. We tested these animals across the commonly used developmental window of 7-28 dpf utilizing a set of light or vibration stimuli to measure non-associative learning (habituation), stress response, fear response, sensory sensitivity and locomotion. Our experiments did not reveal major differences between the two groups, providing a supportive rationale for using the transgenic tg(Elavl3:H2B-GCaMP6s) line for investigating the neuronal circuits regulating behavior. Nevertheless, some minor differences did emerge from our analysis, suggesting that care should be taken in interpreting behavioral assays with mutant/transgenic lines. By performing these behavioral assays at multiple developmental time points, we describe for the first time a longitudinal comparison of responses elicited by the same set of stimuli during larval development.

## 2. Methods

All experiments described in this study involving the use of animals were approved and performed in compliance with the Comité de Protection des Animaux (CPAUL) of Université Laval.

### 2.1 Animal housing and transgenic zebrafish

Adult zebrafish (Danio rerio) of the wild-type Tüpfel long (WT-TL) line and the transgenic line expressing the calcium sensor GCaMP6s tg(Elavl3:H2B-GCaMP6s) in the *casper* mutant were maintained in the fish facility at 28−29 °C with a 14/10 h light/darkness cycle.

Embryos were raised in Embryo medium (NaCl 13.7mM, KCl 0.54mM, MgSO_4_ 1.0mM, CaCl_2_ 1.3mM, Na_2_HPO_4_ 0.025mM, KH_2_PO_4_ 0.044mM, NaHCO_3_ 4.2mM, pH 7.2 with HCl 12N) at 28 °C on a standard 14/10 h on/off light cycle in 100mm x 25mm petri dishes at a ratio of 2mL medium/larva. At 3 days post fertilization (dpf), chorion debris were removed. The medium was renewed from 5 to 10 dpf, and the larvae were fed daily with 30 μL/larva of eukaryotic *Tetrahymena thermophila. T. thermophila* was cultured at 28.5°C in a sterile SPP medium enriched with peptone protease and Fe3+ (2% proteose peptone, 0.1% yeast extract, 0.2% glucose, 33 μM FeCl3, 250 μg/mL Streptomycin/penicillin, 0.25μg/mL Amphotericin B) until their concentration reached a plateau. An 11 ml SPP medium volume was inoculated with 1 ml of *T. thermophila* culture at the maximum confluence. Before use, *T. thermophila* were washed from their SPP medium to remove antibiotics, using three centrifugations (2min, 0.8g) and pellet re-suspension in embryo medium.

At 10dpf, larvae were then transferred into a 1.1L recirculating tank supplied with deionized water supplemented with Instant Ocean® Sea Salt (Tropic Marin, Wartenberg, Germany) and NaHCO_3_ to reach a conductivity between 700-1000 μS and a pH range of 7.2–7.4. Larvae were housed in groups of 25-30 individuals. Water temperature, conductivity, and ammonia levels were recorded daily. The fish were fed daily with 24h decapsulated brine shrimp nauplii (*Artemia nauplii*) until the end of the experiment. The amount of food given was set to be almost entirely eaten in the first 10 minutes to ensure it was not in limited quantity.

### 2.2 Larvae growth

We quantified the growth of both zebrafish lines by measuring standard length (SL) every two days, from 7 to 21 dpf. Larvae were placed on a microscope slide in a drop of solution (either embryonic medium or home tank water) and imaged live with the use of a stereomicroscope (Zeiss Stereo Discovery.V8, 1x objective), a camera (AxioCam MRc; Zeiss) and its associated image acquisition software (AxioVision; Zeiss). Standard length (SL), measured from the snout to the base of the tail, was measured directly from images calibrated to scale with ImageJ (NIH). SL was averaged over three measurements to account for measurement variability.

#### 2.3 Zebrafish Behavioral Assay

All experiments were conducted between 9 a.m. and 5 p.m. Fish were transferred to the experimental room 30 min before the start of the experiment and kept in a 28°C incubator. Behavioral assays for 7 dpf larvae were performed in 24-well plates with 1 larva per well in 1 mL of embryo medium; for ≥ 11 dpf larvae, water from their home tank was used instead. After transferring the larvae to the experimental plate, they were placed on a 29 °C hot plate for 10 min before starting the experiment. The plate was then loaded into a ZebraBox (ViewPoint) containing a computer-controlled lightbox and a video camera with an infrared filter. For each session, data were acquired at 30 fps. The same larva was tested at 7, 11, 14, 21 and 28 dpf, with 12 animals per batch and a minimum of 2 batches.

We used three different protocols:

#### 2.3.1 Light-dark transitions assay (LDA)

The light-dark protocol consisted of three cycles of five minutes of darkness followed by five minutes of white light (set at 9000 lux).

#### 2.3.2 Multi-stimuli assay (MSA)

The multi-stimuli assay consisted of a precise combination of light and vibration stimuli in the following sequence: 5 min of darkness; 5 min of full-spectrum white light (18800 lux); 6 min of strobe light sequence consisting of 3 cycles of 1 min darkness followed by 1min strobe light (10 Hz); 5 min of darkness; 5min of full-spectrum white light. After this sequence, the rest of the protocol was executed in the dark according to the following sequence of stimuli: 84 cycles of 20ms vibrations (400 Hz) with 1s in between each vibration; 3 cycles of 1s light flashes with various light intensity; 4 cycles of 10s light flashes at various light intensities (18800, 2650, 1190, 350 lux); 20ms single vibrations (200 Hz, 400 Hz and 600 Hz); 90 cycles of 20ms vibrations (600 Hz) with 1s in between each vibration.

Figure S1 illustrates the precise experimental sequence. The LDA and MSA were performed using the same animals, which were placed on a warm plate set at 29 °C for 10 min between each protocol.

#### 2.3.3 Light-dark preference

To measure light-dark preference, we used black optic tape to cover half of a 6-well plate (lids and bottom) to separate the wells in half. The larva had the choice to swim to either zone. Larvae were transferred individually in the plate (one per well) and placed on the 29°C hot plate for 10 min before starting the experiment. We then moved the plate to the *ZebraBox* to start the protocol. The experimental condition consisted of 10 min of full-spectrum white light followed by 3 min of strobe light (1 minute of darkness followed by 1 minute of strobe light at 10 Hz) and the final 10 min of full-spectrum white light. A custom algorithm was written in Python to track the larvae centroid in the light area.

### 2.4 Adapted Quantization method

Behavioral characterization was measured using a pixel-difference-based movement detection method (quantization) from the *Zebralab* software (Viewpoint, France). We applied a correction factor to visualize and compare the quantization data from fish with difference in i) size due to the various developmental stages and ii) pigmentation in the GCaMP6s line due to their lack of melanophores, thus being differently processed by the software. To determine the correction factor for each developmental stage, we calculated the mean of movement during the first 5 minutes of the LDA protocol. Then, we set an empirical value to make the mean movement of each stage to correspond to the movement in 28 dpf animals for each line. Once we identified the proper correction factor for each condition, it was applied to the raw pixel difference for all samples. The factor is indicated in the figure legends. We used the same factor to compare the different development stages in wild-type and GCaMP6s lines for consistency.

### 2.5 Locomotion measurements

Larval locomotor activity was quantified using the automated video tracking mode of *ZebraLab* software, which monitored the fish’s position by tracking a centroid on its body. ZebraLab detection parameters were set empirically to detect larval movements with minimum noise. Due to their lack of pigmentation, detection parameters had to be adapted for the GCaMP6s fish, as shown in Table 1. The background was automatically refreshed every second to prevent the appearance of detection artifacts.

**Table 1:** Set of parameters used for the detection of fish with the tracking mode of Zebralab software. X-min size parameters are defined in Zebralab software as the “minimum horizontal size for the object to be analyzed.”

| Days post-fertilization (dpf) | Low Detection Threshold | X Min Size |
| --- | --- | --- |
| 7 | 22 (both lines) | 5 (both lines) |
| 14 | 25 (GCaMP6s fish) / 28 (WT-TL fish) | 6 (GCaMP6s fish) / 7 (WT-TL fish) |

### 2.6 Statistics and data analysis

The raw data generated by the Zebrabox were analyzed using a custom-R script. R graphic programming was used to generate the plots. *P* value computed by Tukey’s honestly significant difference test on 1-way ANOVA. All box plots were generated using R graphic programming and the *ggplot* module. The lower and upper hinges of boxplots correspond to the first and third quartiles. A line indicates the group median. The whiskers extend from the hinges to the maximum or minimum value at most 1.5x the interquartile range (IQR) from the hinge. A *P* value of less than 0.05 was considered significant. The linear regressions were performed using the LM method in the R environment.

## 3. Results

### 3.1 Zebrafish behavioral assays

As they develop, zebrafish undergo behavioral complexification and maturation[4]. It has been shown that zebrafish larvae display stereotypical swimming responses in reaction to various stimuli, including light flashes and acoustic stimulation. We used two complementary assays to investigate the longitudinal evolution of behavioral responses in developing zebrafish larvae: 1) a well-established light-dark transition assay (LDA) consisting of alternating dark-light phases to measure response to light transitions, and 2) an updated version of a previously published protocol[26] which includes a more diverse and complex sequence of stimuli triggering different behavioral responses. This multi-stimuli assay (MSA) consists of both light and vibration stimuli previously associated with fear (strobe light)[27], stress (dark-light transitions)[28], locomotion, sensory sensitivity (light flash and vibration), and non-associative learning (habituation to repeating stimuli)[29]. Notably, these stimuli are commonly used in both behavioral and functional neuronal imaging studies[30]. In both assays, we used changes in locomotion as a readout of behavioral responses.

Our first step was to determine if our assays could replicate the behavior of 7 dpf WT larvae as previously described[26]. We found that the WT larvae exhibited increased movement during the dark phases compared to the light phases (Figure 1A, Figure S1A-S1C). Exposure to strobe light induced a rapid freezing behavior (Figure S1D), followed by a gradual recovery of motor activity. Similarly, single light flashes led to a rapid decrease in movement, akin to freezing (Figure S1E). We also observed habituation to the fast repetition of vibration bursts (Figure S1F), while single vibration events elicited a brief escape response (Figure S1G). These findings suggest that our LDA and MSA assays successfully reproduced the behavioral responses previously described in 7 dpf WT larvae[26].

**Figure 1:**
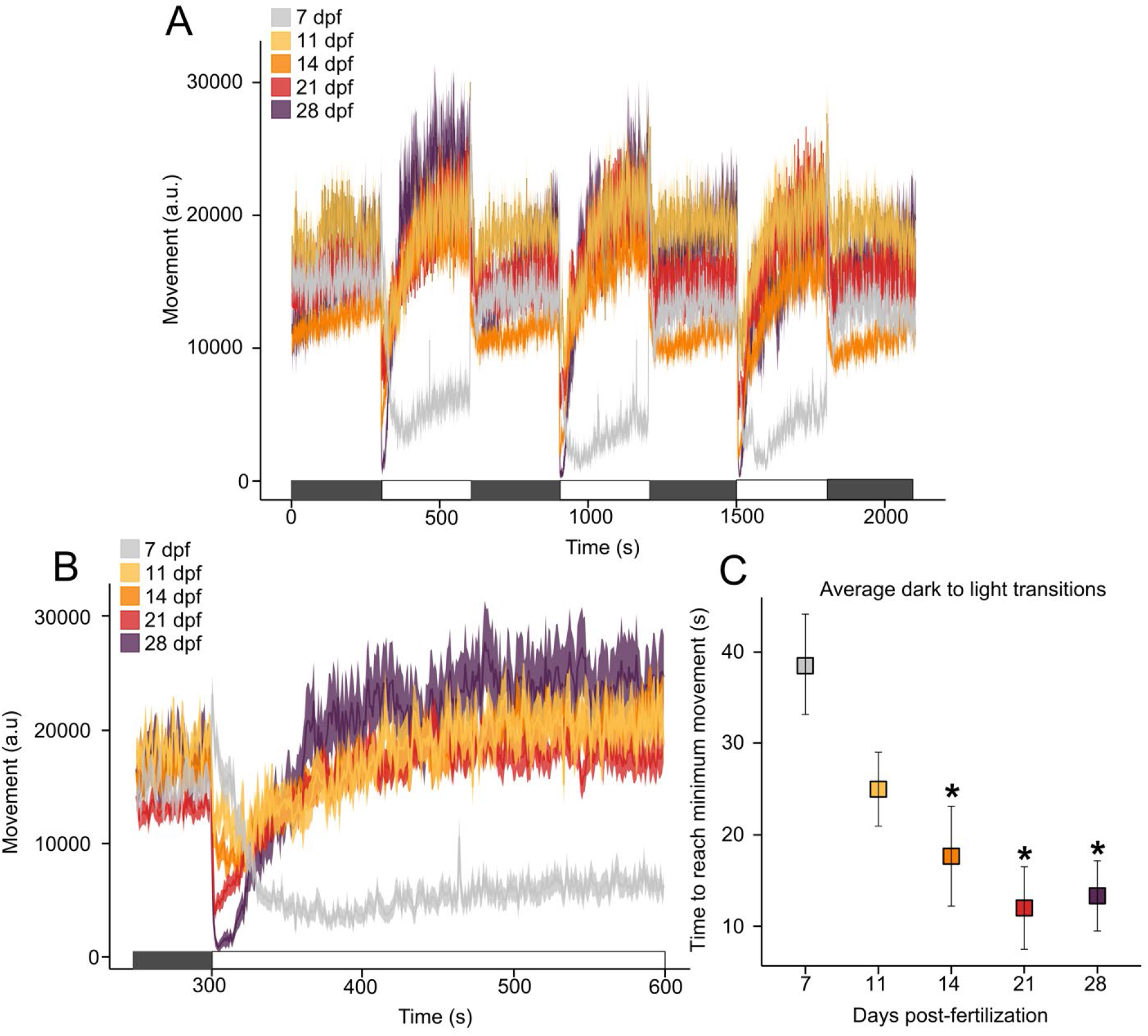
Shift of sensorimotor responses to light-dark transitions in wild-type zebrafish during the second week of larval development. (**A**) Larvae movement during the Light-Dark Assay (LDA) phases across zebrafish larvae development. The data represent the average movement for each condition. (**B**) The transition from dark to light induces a gradual reduction in movement in 7 dpf larvae, whereas from 11 dpf onwards, there is an initial immediate freezing response followed by a rapid locomotion increase. (**C**) The time required to reach minimum movement after transitioning from dark to light decreases significantly across development. Data are shown as the mean ± SEM, n = 48 – 60 for each time point. *P value* compared each time point versus 7 dpf. *p <0.05. *P* value computed by Tukey’s HSD significant difference (Honestly Significant Difference) test on 1-way ANOVA [F(4,10) = 30.81, *P =* 1.34^e^-05]. The correction factors used across development are as follows: 20 (7dpf), 10 (11dpf) and 5 (14dpf). These experiments were performed using a within-subjects design.

### 3.2 Sensorimotor responses across development

We repeated both assays across larval development to establish a behavioral profile over the most commonly studied developmental window used for functional imaging and pharmacological studies: 7 to 28 dpf.

#### 3.2.1 Responses to light stimuli

An intriguing feature that differentiates zebrafish larvae from adults is their light-dark preference, with larvae exhibiting a preference for light[31] in contrast to the preference for darkness observed in adults[32]. We wondered whether the response to light transitions also changes during development and, if so, when. Thus, we compared 1) the overall activity in both dark and light phases and 2) the responses to visual transitions at different time points during larval development.

In 7 dpf WT larvae, the dark-to-light transitions triggered an initial burst of movement that gradually decayed, reaching a persistent freezing state throughout the light phase (Figure 1B-C and Figure 2A-B). A similar burst was observed in light-to-dark transitions, while overall activity remained high during the dark phases (Figure 2C and Figure S2A). In contrast, starting at 11 dpf and onward, the larvae’s initial reaction to dark-to-light transitions was a sharp and transient reduction in movement, reaching complete freezing at later developmental stages, followed by a gradual increase in activity (Figures 1B-C and 2B). Meanwhile, the light-to-dark transition did not elicit a burst of movement in larvae older than 7 dpf (Figures 2C and S2).

**Figure 2:**
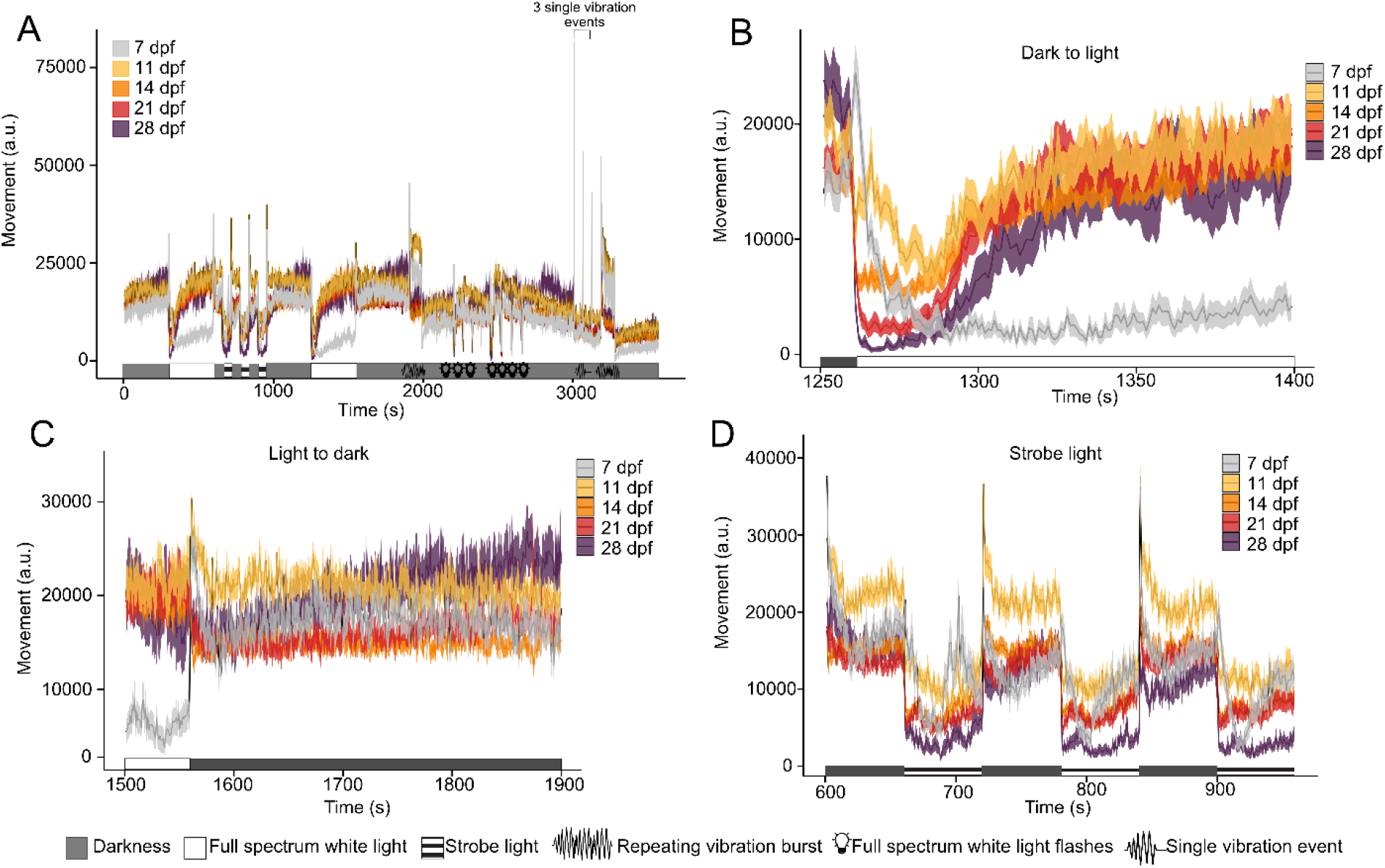
Sensorimotor responses to Multi-Stimuli Assay in wild-type zebrafish. (**A**) Movement responses to Multi-Stimuli Assay (MSA) across zebrafish development. The data represent the average movement for each condition. (**B**) Following a dark-to-light transition, 7 dpf larvae show an initial burst of activity followed by a rapid decrease in movement, which persists for the duration of the light phase. Beyond 7 dpf, the transition to light induces an initial freezing response, followed by a subsequent return to baseline activity. (**C**) The transition from light to dark induces a peak activity only in 7dpf WT-TL larvae. (**D**) Stroboscopic light induces a sharp reduction in movement; after a few seconds, larvae start moving again in all stages. Data are shown as the mean ± SEM, n = 24-36 for each time point. The correction factors used across development are as follows: 20 (7dpf), 10 (11dpf) and 5 (14dpf). These experiments were performed using a within-subjects design.

During strobe-light stimulation, we observed a consistent freezing behavior across larval development (Figure 2D). Moreover, as larvae develop, we observed a gradual reduction in total movement during strobe light stimulation (Figure S3A). The reduction in movement is followed by a progressive recovery of activity at all developmental stages, which is more pronounced in 7 dpf larvae. (Figure 2D et S3B).

Overall, unlike older animals, 7 dpf larvae exhibited consistently higher activity levels during dark phases than light ones (Figure S4). Taken together, the main difference we observed was a change in light-dark transitions that may reflect a development milestone appearing between 7 dpf and 11 dpf. We also observed changes in their responses to strobe light stimuli that could reflect a different sensitivity to changes in light conditions.

#### 3.2.2 Responses to vibrations

To expand our characterization beyond light stimulation, we exposed larvae to different sequences of short bursts of vibration, which triggers a sudden increased motor activity (Figure 3A-B and S1F-G). This behavior was observed in all behavioral stages (Figure 3A). We first detected a greater initial response in 7 dpf larvae compared to other developmental stages, while the lowest response was observed in 14 dpf larvae (Figure 3A-B). As previously described[29], larvae showed habituation to repeated short vibration bursts (Figure 3A-C, Figure S1F), and 7 dpf animals showed the most rapid reduction in responses across time.

**Figure 3:**
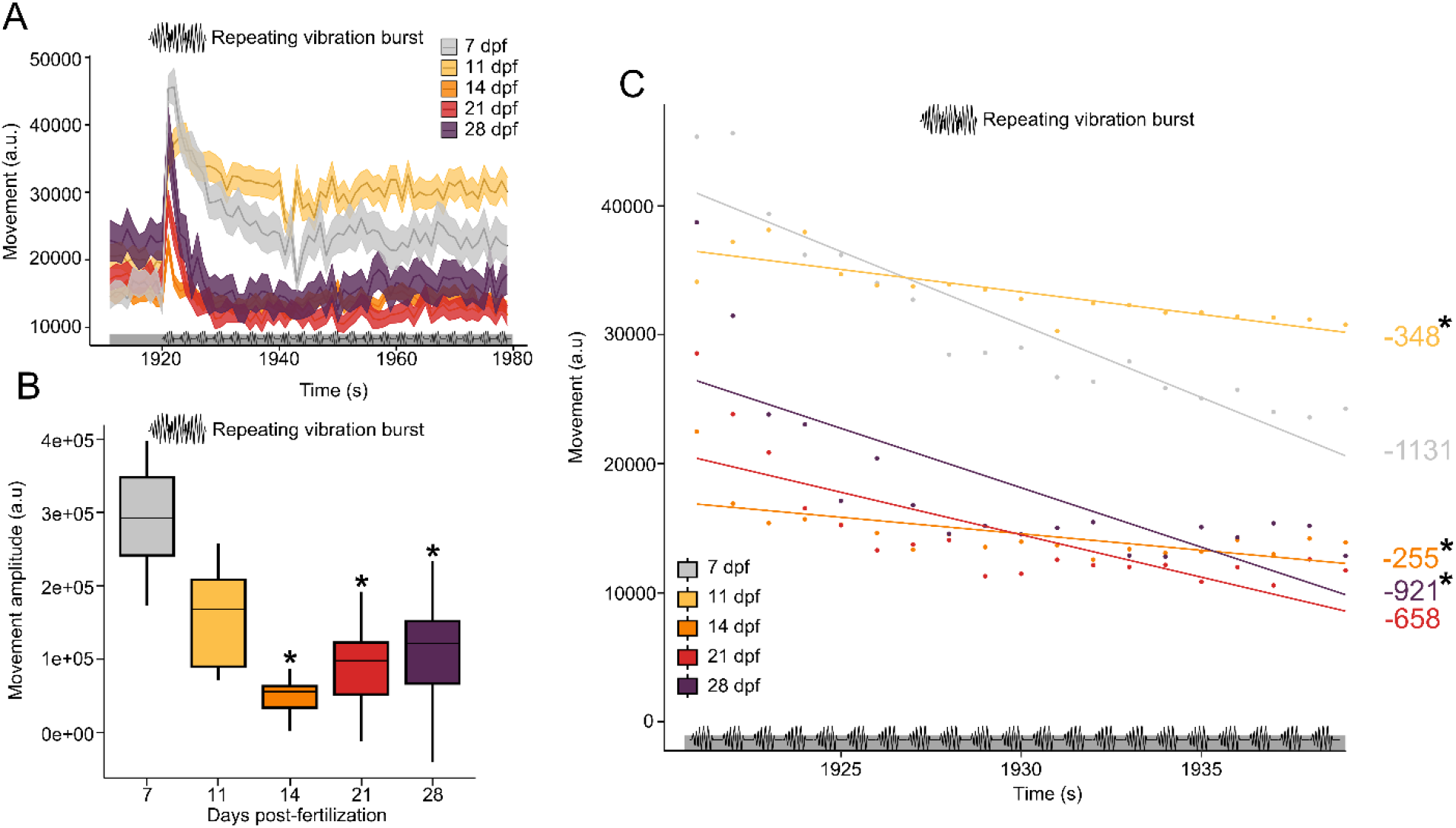
Developmental U-shaped sensorimotor responses to vibratory stimuli in the Multi-Stimuli Assay in WT-TL zebrafish. (**A**) Repeating short burst of vibration (400Hz) induces an initial rapid response followed by a gradual habituation to the stimuli. (**B**) Amplitude of movement after the first vibration burst (400Hz) of the habituation sequence. The amplitude is calculated by comparing the response to the vibration to the average of movement during the 3 seconds immediately before the stimuli. n = 24-36 for each time point. The correction factors used across development are as follows: 20 (7 dpf), 10 (11 dpf) and 5 (14 dpf). *P* value computed by Tukey’s HSD significant difference test on 1-way ANOVA [F(4,55) = 12.12, *P =* 3.94^e^-07]. (**C**) Linear regression of fish movement following repeating vibration burst shows habituation to this repeating stimulus. The slope for each regression shows that 7 dpf animals display a more rapid habituation to repeating stimuli. *P* value computed by Tukey’s HSD significant difference test on 1-way ANOVA [F(4,55) = 10.64, *P =* 1.86^e^-06]. *p <0.05 when compared to 7 dpf. These experiments were performed using a within-subjects design.

As opposed to light stimuli, we observed consistent behavioral responses to vibration burst but the intensity of the responses is modulated across development with 7 dpf larvae showing the greater response.

### 3.3 Behavioral profile of the Elavl3:H2B-GCaMP6s line

The transgenic line used in our study has been developed to measure neuronal activity through the expression of a nuclear calcium sensor, GCaMP6s, under the control of a pan-neuronal promoter (HUC/elavl3) tg(Elavl3:H2B-GCaMP6s) in the *casper* mutant. These fish are pigment-free and, therefore, highly transparent due to the mutations of two of the three genes coding for skin pigments in zebrafish (*nacre* and *roy*)[13].

#### 3.3.1 Growth and movement comparison

To compare the sensorimotor responses of the GCaMP6s *versus* WT-TL lines over development, we first analyzed their growth by measuring their length every 2 days from 7 to 21 dpf. From the second week of development, we observed that GCaMP6s animals were slightly, but significantly smaller than WT animals. (Figure S5).

To investigate locomotion more precisely, we measured additional parameters, such as distance travelled, using the tracking feature of *ZebraLab* software. Using data from the LDA, we detected an increase in overall distance travelled up to 14 dpf in both lines and a non-significant decrease in locomotion in the transgenic line compared to the wild-type from 7 dpf to 21 dpf (Figure S6).

#### 3.3.2 Sensorimotor responses comparison between lines

To determine whether sensorimotor responses were altered in the GCaMP6s line, we repeated the behavioral profiling using the light-dark and multi-stimuli assays at the same developmental stages as previously performed. Overall, in the GCaMP6s line, we also observed developmental differences in responses to visual and vibration stimuli as with the WT-TL line (Figures S7-S8-S9-S10). In the GCaMP6s line, we also observed a switch in responses to dark-light transitions (Figures S7A-7C and S8B-8C) and an increase sensitivity to vibrations in 7 dpf larvae (Figure S9). However, unlike WT-TL, GCaMP6s larvae of 14 dpf and older did not show significantly higher activity during light phases. (Figure S10).

As we performed the behavioral characterisation under the same conditions in both lines, we then sought to compare the sensorimotor responses of GCaMP6s and WT-TL lines directly.

Although the overall behavioral profiles of the GCaMP6s and WT-TL lines were similar across developmental stages in the LDA and MSA assays (Figures 4 and 5), we detected some differences. At 7dpf, we did not detect any differences in responses to light-dark transitions between WT-TL and GCaMP6s fish (Figure 6A). However, during the strobe light phases, the GCaMP6s line exhibited a more persistent freezing behavior, as reflected by a reduction in overall movement and a greater freezing index (Figure 6C-D). Regarding the response to vibration stimuli, 7 dpf GCaMP6s larvae demonstrated a higher initial response but had similar habituation to WT-TL larvae in response to repeating vibration bursts (Figure 6B). Similar differences were also measured at 14 and 21 dpf where GCaMP6s animals had a higher reduction in total movement during the strobe light than WT (Figure S11) and an increased response to vibration burst (Figure 7A).

**Figure 4:**
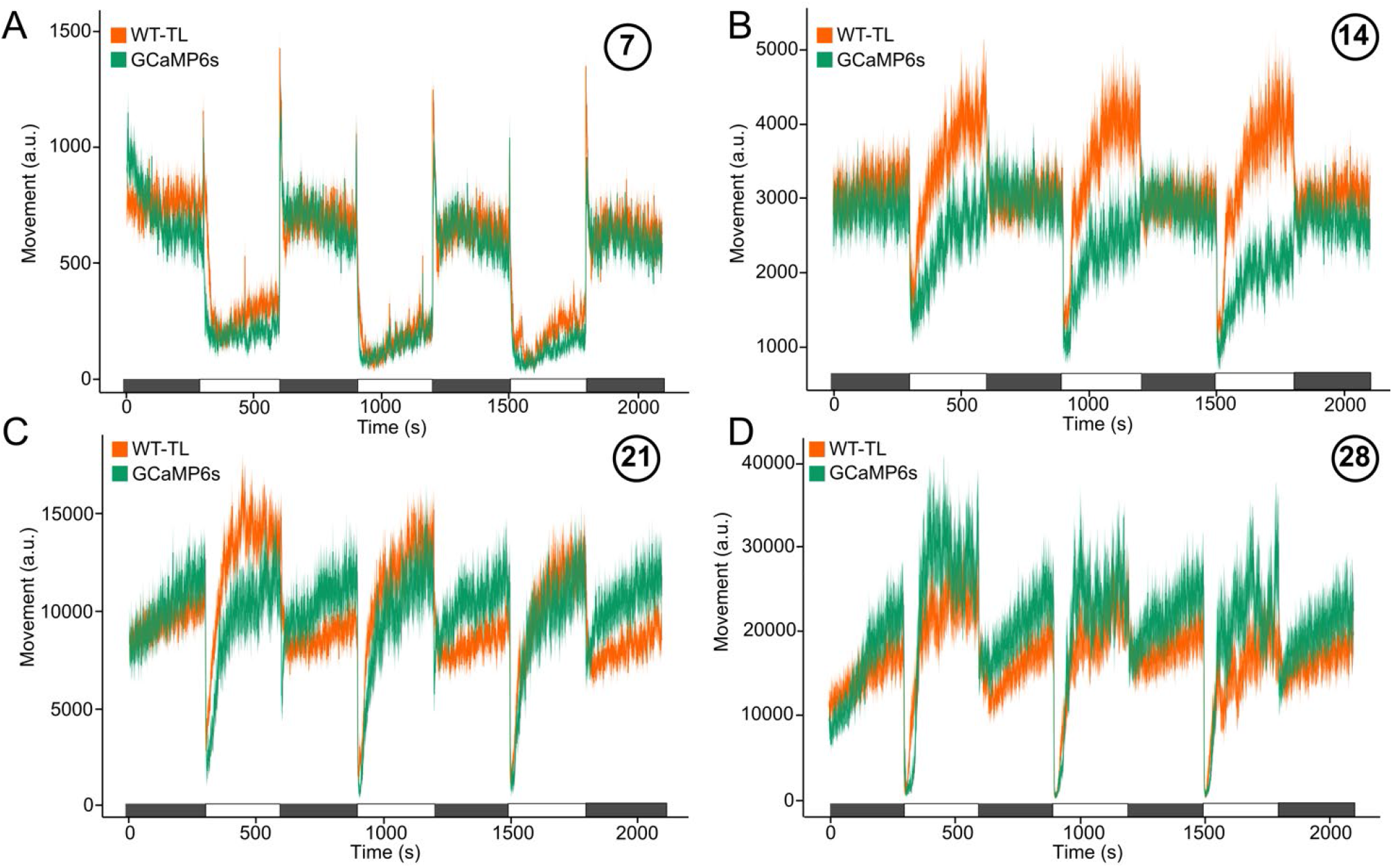
GCaMP6s zebrafish larvae exhibit similar sensorimotor movement response to light-dark transitions, compared to WT-TL, albeit with different relative amplitudes. Movement of the fish during Light-Dark-Assay at (**A**) 7, (**B**) 14, (**C**) 21, and (**D**) 28 dpf for WT-TL and GCaMP6s lines, plotted in 1s time bins. Data are shown as the mean ± SEM, n = 48 – 60 animals for both lines and for each time point. The correction factors used for transgenic animals are 2 (7 and 28 dpf) and 3 (14 and 21dpf). These experiments were performed using a between-subjects design.

**Figure 5:**
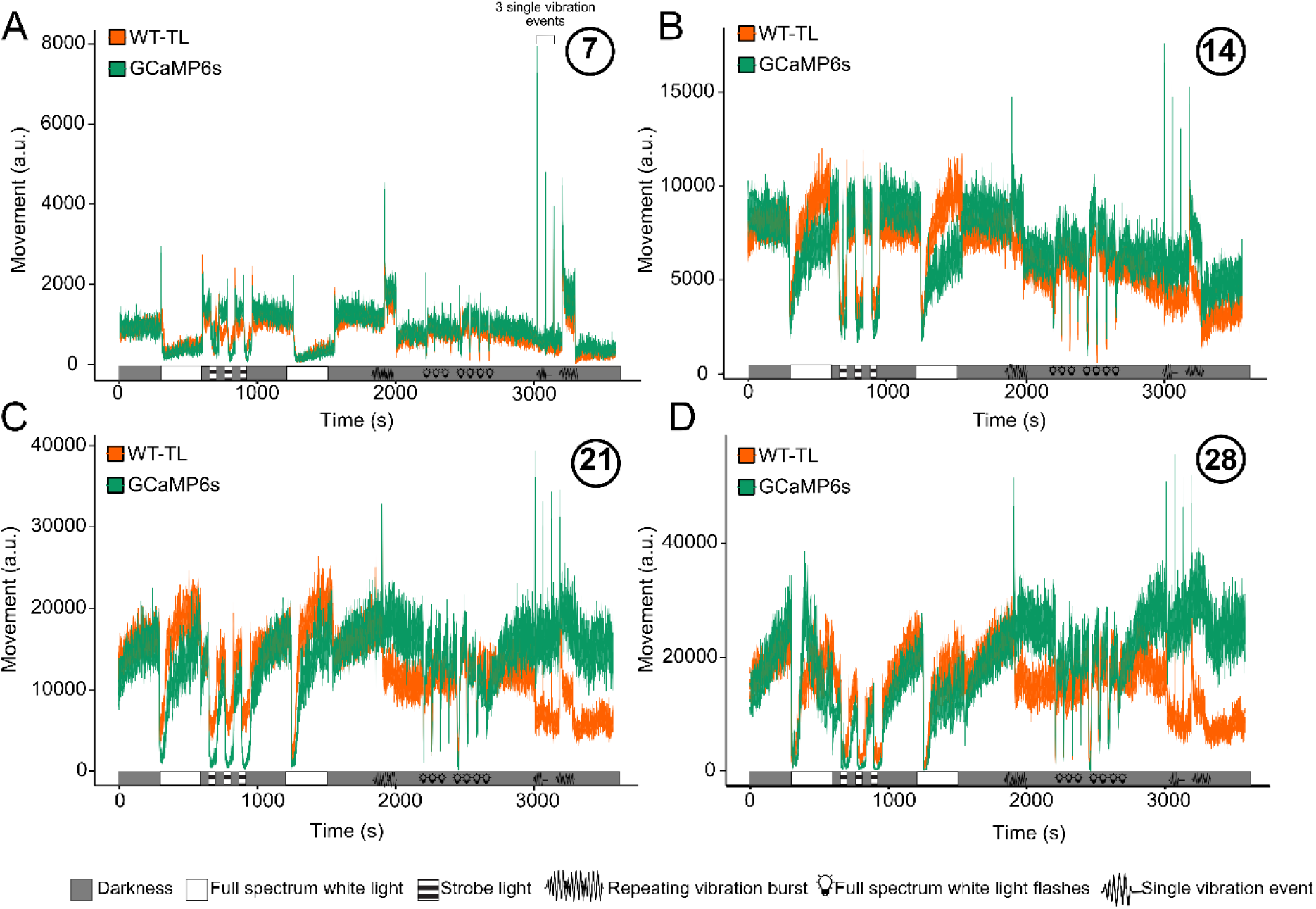
Sensorimotor responses by WT-TL and GCaMP6s lines during Multi-Stimuli Assay across development. Total movement of WT-TL and GCaMP6s fish at **(A)** 7, **(B)** 14, **(C)** 21, and **(D)** 28 dpf plotted in 1s time bins. Data are shown as the mean ± SEM, n = 24 – 36 for both lines and for each time point. The correction factors used for GCaMP6s animals are 2 (7 and 28 dpf) and 3 (14 and 21dpf). These experiments were performed using a between-subjects design.

**Figure 6:**
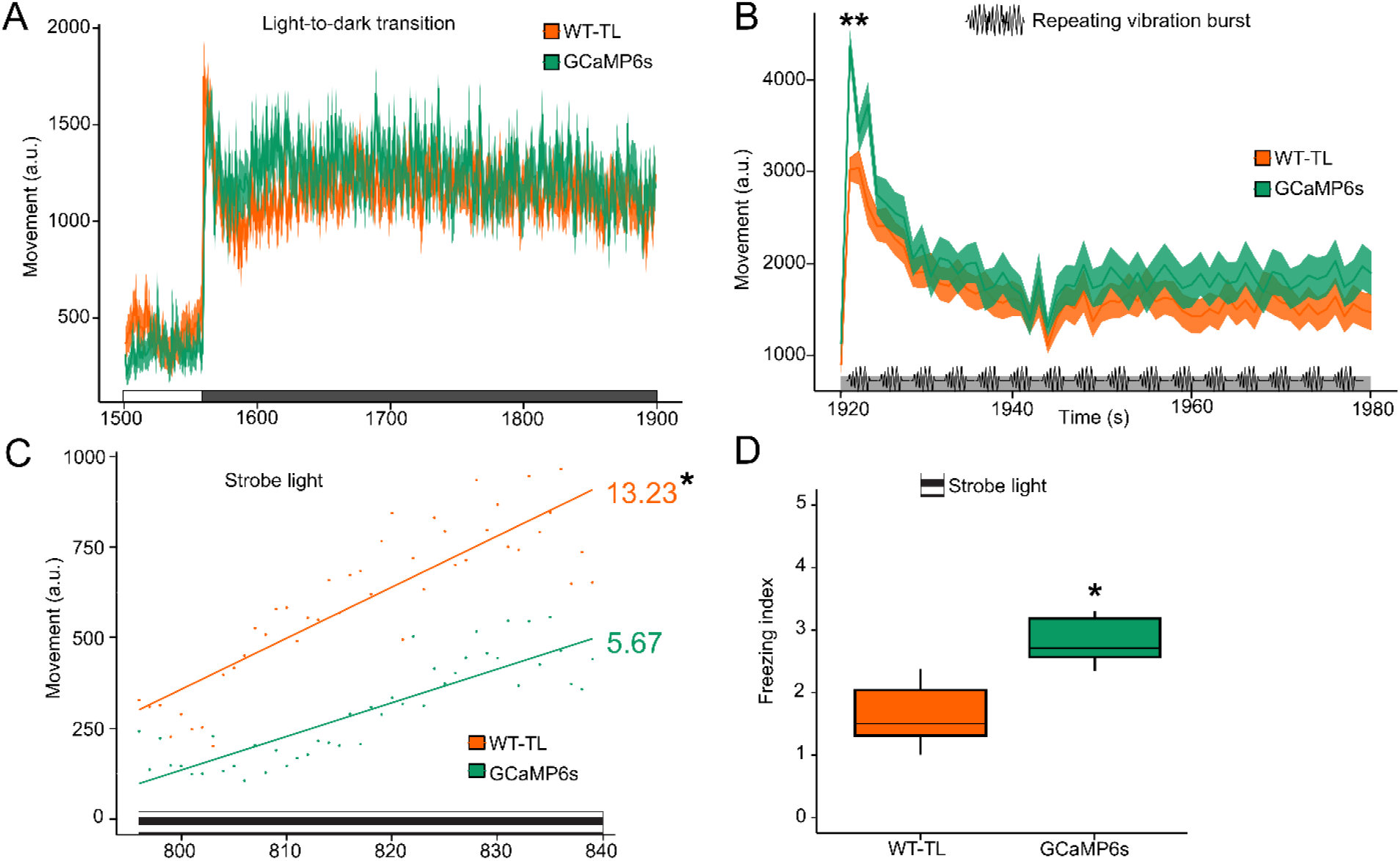
Strobe light stimuli elicit a lower motor activity response in the GCaMP6s line than WT-TL at 7 dpf. (**A**) Wild-type and GCaMP6s larvae show no difference in movement activity during light-to-dark transitions. (**B**) GCaMP6s larvae exhibit a greater initial response to repeating vibration bursts than WT-TL. Both lines demonstrate similar habituation to repeating stimuli. *P* value computed by Tukey’s HSD significant difference test on 1-way ANOVA [F(1,355) = 13.58, *P =* 0.0002] (**C**) GCaMP6s larvae display significantly slower habituation during the strobe light compared to WT-TL. *P* value computed by Tukey’s HSD significant difference test on 1-way ANOVA [F(1,355) = 13.58, *P =* 0.0002] (**D**) The freezing index indicates the ratio of the sum of movements during dark phases over the sum of movements during strobe light phases. GCaMP6s larvae have a greater freezing index than WT-TL, indicating reduced activity during strobe light. *P* value computed by Tukey’s HSD significant difference test on 1-way ANOVA [F(1,22) = 30.83, *P =* 1.4^e^-05]. n = 36 for both lines. *\*p*<0.05, **p<0.01. The correction factors used for transgenic animals at 7dpf is 2. These experiments were performed using a between-subjects design.

**Figure 7:**
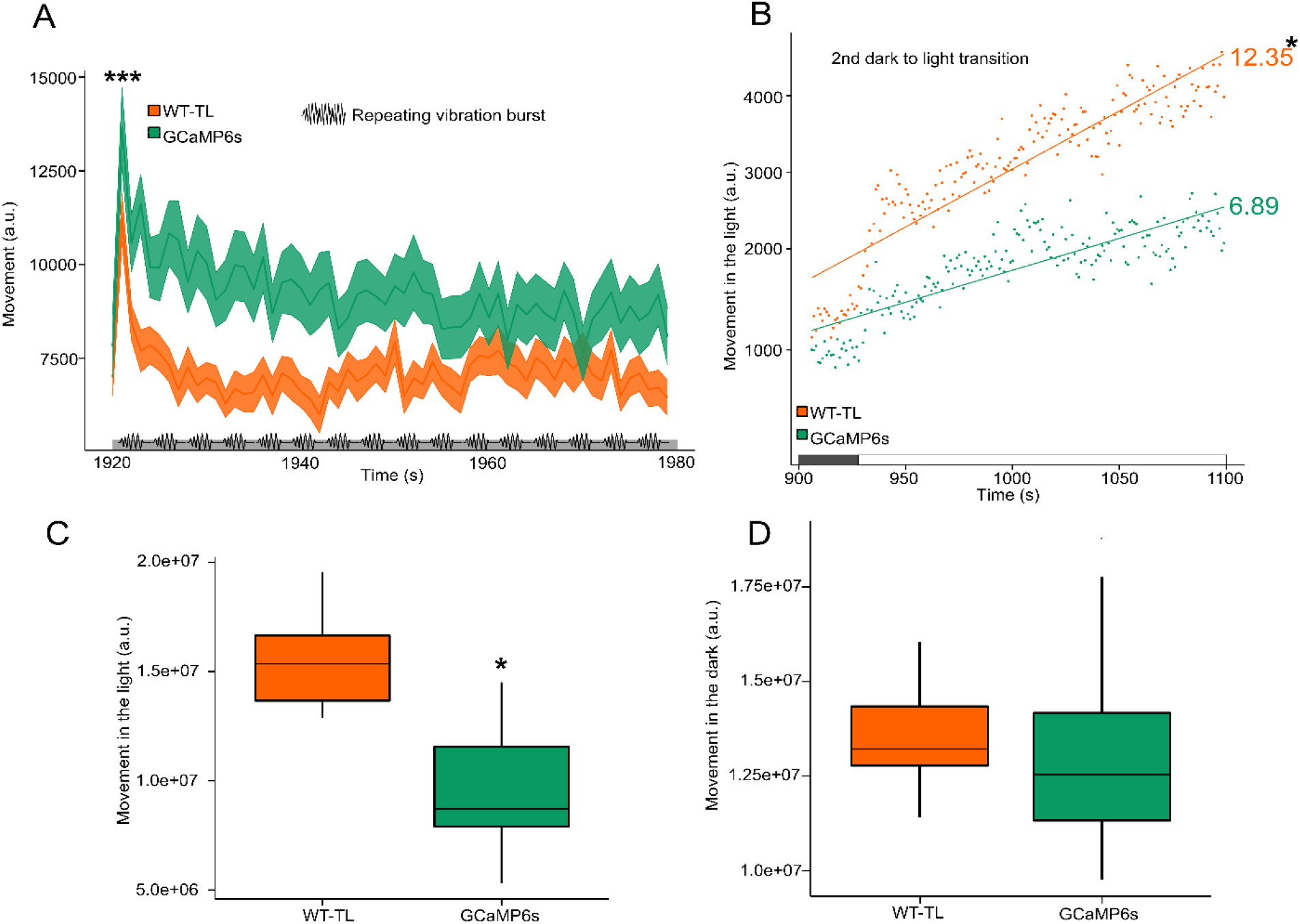
GCaMP6s larvae display higher sensitivity to repeating vibration stimuli but respond less to dark-to-light transitions at 14 dpf than WT-TL. (**A**) GCaMP6s larvae exhibit a greater initial response and slower habituation to a repeating vibration burst than WT-TL larvae. *P* value computed by Tukey’s HSD significant difference test on 1-way ANOVA [F(1,117) = 12.96, *P =* 0.000467]. (**B**) Linear regression of fish movement following the second transition from dark to light. GCaMP6s larvae exhibit a significantly slower recovery rate following the initial freezing response. *P* value computed by Tukey’s HSD significant difference test on 1-way ANOVA [F(1,22) = 11.3, *P =* 0.003]. (**C**) GCaMP6s fish have decreased total movement during light phases *P* value computed by Tukey’s HSD significant difference test on 1-way ANOVA [F(1,22) = 33.91, *P =* 7.37^e^-06]. (**D**) but show no difference in activity during dark phases compared to WT-TL larvae. *P* value computed by Tukey’s HSD significant difference test on 1-way ANOVA [F(1,22) = 0.244, *P =* 0.626]. n = 60 for both lines. *\*p*<0.05, ***p<0.001. The correction factors used for transgenic animals at 14 dpf is 3. These experiments were performed using a between-subjects design.

Moreover, at 14 dpf, we also observed a delayed recovery following a dark-to-light transition (Figure 7B) and reduced total movement only during light phases (Figure 7C-D). By 21 dpf, we detected a difference only in the recovery from strobe light phases (Figure S12), and by 28 dpf, both strains had almost identical responses.

Interestingly, during the MSA at later developmental stages (14, 21 and 28 dpf), the GCaMP6s fish exhibited an increase in locomotion, which began after the initial step of the habituation phase (repeating vibration burst) (Figure 5C-D). Despite this hyperactivity, we did not measure differences in habituation rate, sensitivity to light stimuli or vibration at any developmental stage between the WT-TL and GCaMP6s lines. To determine if this hyperactivity was specific to the end of the MSA, we calculated a movement index at both ends of the protocol. The WT-TL line showed reduced movement at the end of the protocol (Figure S13 A-D), whereas the GCaMP6s line moved more at the end at 14, 21 and 28 dpf (Figure S13 C-D).

Overall, we observed similar behavioral responses to the different stimuli in both lines, albeit sometimes with different amplitude or duration.

#### 3.3.3 Light-dark preference

We observed a persistent freezing behavior in the GCaMP6s line during light transitions (Figures 4, 5, 6 and S8) which might suggest increased anxiety levels compared to the WT-TL line[31]. Light-dark preference assays have been used to assess anxiety-like behavior in zebrafish, where a fish can swim between covered (dark) and uncovered (light) area[31–33]. We selected two representative developmental stages, 7 and 14 dpf, to investigate anxiety-like behavior in both lines. We measured the time spent in the light zone during the first 10 min of the experiment, then applied a burst of strobe light as an aversive stimulus, and finally measured the light-dark preference again (Figure S14A). Figures S14B and S14C illustrate that both lines spent a similar amount of time in the light zone during the three periods of the experiment at 7 and 14 dpf. However, a slight difference (p = 0.0782) was detected in the time spent in the light zone during the final 10 minutes of the experiment at 14 dpf, with GCaMP6s larvae spending less time in the light zone compared to WT-TL larvae (Figure S14C). Taken together, the result from this test doesn’t support an increased baseline stress level between the GCaMP6s and the WT-TL larvae.

## 4. Discussion

This study describes a detailed description of the evolution of sensorimotor responses by larval zebrafish during a well-studied developmental period in this model. We observed a developmental switch in visual-motor response between 7 and 11 dpf in both WT-TL and GCaMP6s lines. Additionally, we noted a similar overall pattern of sensorimotor responses by a zebrafish line (tg(Elavl3:H2B-GCaMP6s)) that is widely used in studies investigating the relationship between brain activity and behavior[18, 19, 34]. Firstly, we wanted to compare the behavioral responses of zebrafish larvae during their most studied developmental period. To achieve this, we conducted the same behavioral tests every seven days. Secondly, we aimed to verify that the pan-neuronal expression of the calcium sensor GCaMP6s in the *casper* line does not impact the behavioral response of these fish at any point during their early developmental stages.

### 4.1 Sensorimotor responses over larval development

As animals grow and develop, their behavior becomes more complex. This is true across species, whether the animal is learning to communicate, move, or interact with other conspecifics. Zebrafish larvae are particularly well suited to study the neuronal processes driving such complexification. For instance, during the first month after fertilization, larvae develop social preference, hunting instinct, and associative learning abilities[6–8, 35]. Hence, this developmental window is commonly used for functional brain imaging and pharmacological studies investigating the neurobiological mechanisms regulating behavior. Recognizing the significance of this critical period, we decided to compare sensorimotor responses over the first 28 days post-fertilization to further understand the early behavioral maturation of larval zebrafish.

While overall, behavioral responses such as habituation, freezing behavior and reaction to sensory stimuli for each line remained similar across development, we detected a significant difference in responses to dark/light transitions and total movement during these phases as early as 11 dpf. While young larvae (7 dpf) tended to freeze under light, more mature ones (> 11 dpf) exhibited increased activity. Furthermore, while the transition from dark to light triggered a rapid burst in activity in young larvae, it led to a decreased activity in older ones (> 11 dpf). Importantly, these differences were observed in both lines. These observations suggest a developmental milestone in zebrafish sensorimotor behavior, which aligns with the fact that larvae and adult zebrafish differ in their preference for light/dark environments. Notably, adult zebrafish are more active during light phases compared to darkness[31, 33].

The consistently greater initial response to changes in sensory stimuli (light transitions, vibration events) observed in younger animals might reflect less elaborated sensorimotor processing at this early developmental stage. Interestingly, a few studies have identified extensive brain remodelling during the second week of life, such as the refinement of the receptive fields of optic tectum neurons[20]. Similarly, different sensory organs, such as the lateral line, which specializes in water movement detection, undergo significant maturation during fish development[36].

The mechanisms driving these differences remain to be identified. Our findings will be instrumental in studying the neurobiology of the sensorimotor response evolution in zebrafish development.

### 4.2 Sensorimotor responses by the GCaMP6s line

Functional neuronal recording in live animals represented an important breakthrough for zebrafish neuroscience research, enabling the study of brain activity at cellular and circuit levels. Transgenic lines expressing a calcium sensor (GCaMP) in a transparent background are commonly used to investigate neuronal networks involved in opto-motor responses, learning paradigms and sensory responses[14, 17, 30]. Our study provides a detailed assessment of the responses by a GCaMP line over the most commonly studied developmental window. Overall, our longitudinal comparison of behavioral responses of a *casper* line expressing Elavl3:H2B-GCaMP6s and a wild-type line did not reveal any major differences, as both groups showed similar responses to light transitions, vibrations, and habituation to repeated stimuli. Interestingly, a previous study found no difference in associative learning using visual cues at the early larval stage between the transgenic line expressing pan-neuronal GCaMP and WT lines[23]. While we obtained similar results, a notable difference between our studies and Yang et al. [23] was the use of different GCaMP6 variants (fast vs slow), which could have impacted calcium homeostasis differently.

The observed delay in the recovery time in GCaMP6s fish following light transitions and the rapid freezing behavior could indicate increased anxiety in these animals. However, in the light-dark preference assay at 14 dpf, we detected only a slight, non-significant decrease in the time spent in the light zone by GCaMP6s larvae. The differences observed might instead result from an increased sensitivity to white light by GCaMP6s fish, as responses to another repetitive stimulus (vibrations) were similar between the two lines.

In addition, the persistent increase in overall activity in GCaMP6 fish following the vibration stimuli in the MSA at later developmental stages (14 dpf, 21dpf and 28 dpf) could support increased sensitivity to sensory stimulation.

Although the pattern responses to most stimuli were similar between the two lines, we nevertheless detected slightly lower overall locomotion in the GCaMP6s line compared to the WT-TL line, as well as differences in responses to light transitions. GCaMP6s line demonstrated a slower recovery in response to light compared to the WT-TL line. We believe that the differences observed in response to light transitions are not only due to hypoactivity, as responses to other stimuli remained unchanged. Additionally, we observed a slightly lower growth rate for the GCaMP6s line, which may be due to several factors, such as reduced hunting efficiency or genetic factors.

Overall, our data provide important insights for developing combined functional imaging studies and behavioral analyses, exploiting GCaMP-expressing lines to assess how changes in functional circuit connectivity influence behavior in developing zebrafish. While this study focused on responses to specific stimuli, investigating more complex behaviors, such as associative memory or novel-object recognition, would offer further understanding of the behavioral complexification of zebrafish larvae as they develop.

## Abbreviations

Tg: transgenic
dpf: days post-fertilization
WT: wild-type
LDA: Light-Dark assay
MSA: Multi-stimuli assay

## Data statement

Data and custom tracking algorithms are available upon request.

## Notes

The authors declare no competing financial interest.

## Authorship

M.C Conceptualization, Methodology, Investigation, Formal analysis, Visualization, Writing – Original draft, Writing – Reviewing and editing.

M.R.C Investigation, Formal analysis.

C.H Software, Formal analysis, Data Curation, Writing – Reviewing and editing.

N.I Software, Formal analysis, Data Curation.

S.P Software, Formal analysis.

M.L Supervision, Methodology, Writing – Reviewing and editing.

P.D.K Conceptualization, Resources, Supervision, Funding acquisition, Writing – Reviewing and editing.

G.D.B Conceptualization, Formal Analysis, Resources, Methodology, Software, Validation, Visualization, Data Curation, Supervision, Funding acquisition, Project Supervision, Writing – Original draft, Writing – Reviewing and editing.

## Acknowledgments

We want to acknowledge the LARSEM and the animal care facility at CERVO for providing Zebrafish husbandry, laboratory space, and equipment to carry out portions of this research. This work was supported by National Science and Engineering Research Council of Canada (P.D.K # RGPIN-2023-05980, G.D.B #RGPIN-2023-05336), by the Sentinel North program of Université Laval (Canada First Research Excellence Fund), and by the Northern Contaminants Program of Canada. Tg(Elavl3:H2B-GCaMP6s) was developed by Misha Ahrens (Janelia, USA) and transferred to our facility from Ed Ruthazer’s lab (Montreal Neurological Institute, McGill, Canada).

