## Supplemental Figures for "Development of sensorimotor responses in larval zebrafish: a comparison between wild-type and GCaMP6s transgenic line"

**Affiliations:** [1]CERVO Brain Research Centre, Québec City, Québec, Canada, [2]Université Laval, Department of Psychiatry and Neurosciences, Faculty of Medicine, Québec City, Québec, Canada [3]Université Laval, Department of Biochemistry, Microbiology and Bioinformatics, Faculty of Science and Engineering, Québec City, Québec, Canada, [4]Department of Neurobiology and Molecular Medicine Program, University of Utah School of Medicine, Salt Lake City, UT, USA.

**Supplemental Figures**

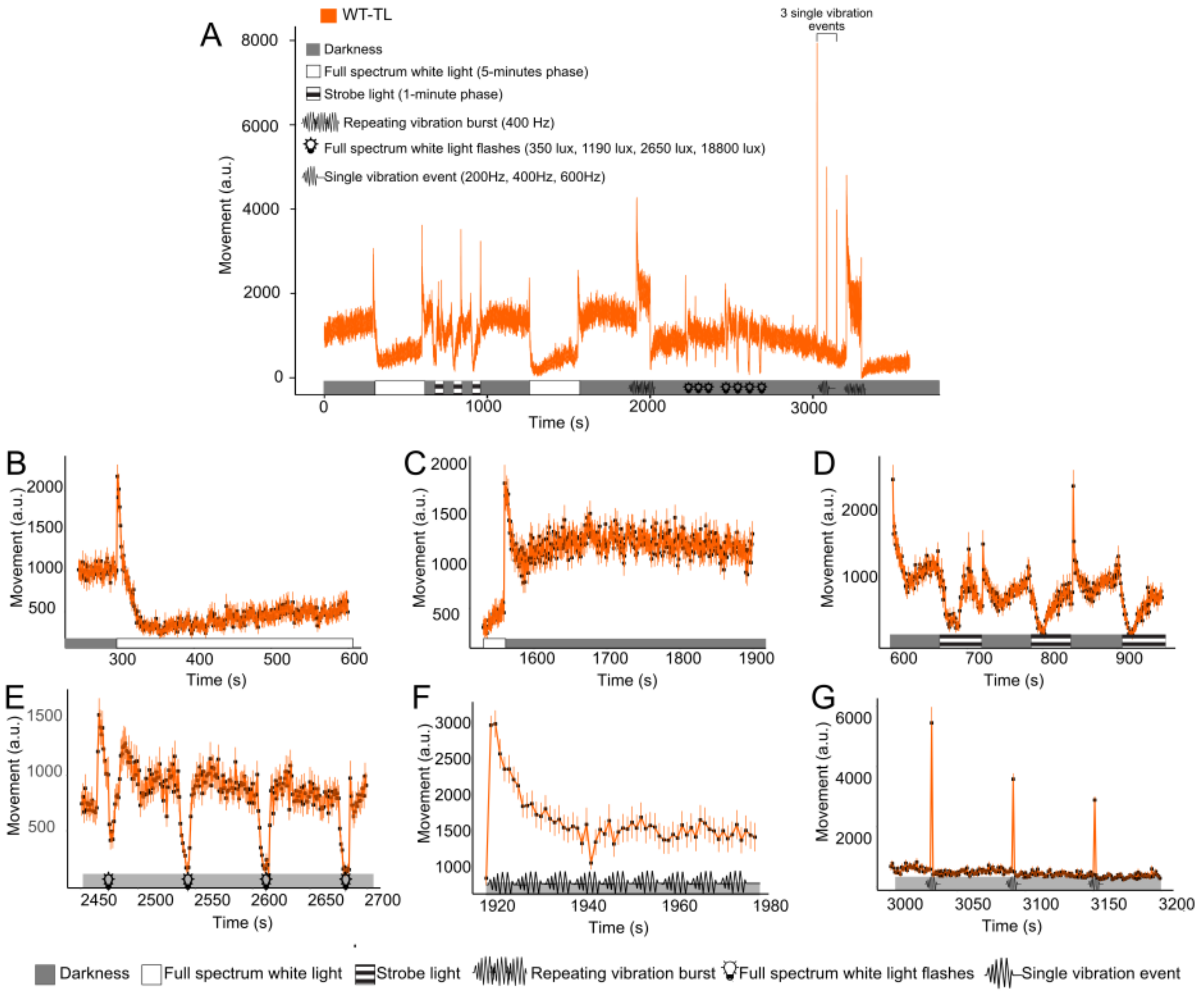


**Figure S1: Validation of the Multi-Stimuli Assay protocol at 7 dpf in wild-type larvae.** (**A**) Overview of the different stimuli performed during the Multi Stimuli Assay (MSA) and the behavioral profile of wild-type (WT-TL) animals at 7 dpf. The different stimuli and their corresponding responses are illustrated. (**B**) The transition from darkness to 100% full spectrum white light induces an initial peak in activity followed by a rapid decrease in movement. Fish then remain calm for this 5-minute phase. (**C**) The transition from 100% full spectrum white light to darkness induces a sharp increase in activity followed by a slow decline in activity. Overall locomotion remains high afterwards. (**D**) Strobe light sections correspond to a succession of 1-minute darkness followed by 1-minute strobe light. The transition induces a very rapid freezing behavior. After a few seconds, larvae start moving again. The transition back to darkness leads to an increase in locomotion. (**E**) Flash of 10 seconds of white light of different intensity (350, 1190, 2650 and 18800 lux). Light flashes lead to a rapid decrease in activity, followed by a rapid return to baseline activity. (**F**) Habituation to a repeating sequence of short vibrations burst (400 Hz). Fish initially react strongly to the vibrations but rapidly habituate and decrease their responsiveness as the sequence progresses. (**G**) Single vibration burst of increasing intensity (200 Hz, 400 Hz, 600 Hz). Fish have a short and consistent burst of activity following the single vibration event, but the amplitude decreases with each stimulus. Each point corresponds to the average movement of all tested animals.

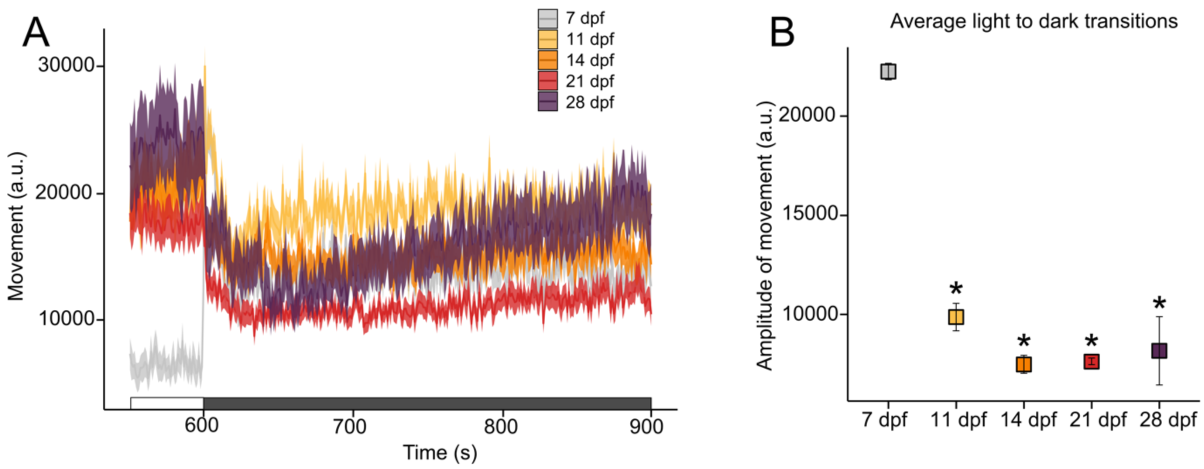


**Figure S2: Light-to-dark transition triggers a transient burst of activity only in 7 dpf WT-TL larvae.** (**A**) Representative traces of sensorimotor response triggered by the light-to-dark transition at each developmental stage. (**B**) Light-to-dark transition elicits a significative change in activity amplitude in 7dpf WT-TL larvae, which is absent at other developmental stages. Data are shown as the mean ± SEM, n = 48 – 60 animals for each time point. *P value* compared each time point versus 7 dpf. *p <0.05. *P* value computed by Tukey's honestly significant difference (Honestly Significant Difference) test on 1-way ANOVA [F(4,10) = 51.79, *P* = $1.2^{e}$-06]. The correction factors used across development are as follows: 20 (7dpf), 10 (11dpf) and 5 (14dpf). These experiments were performed using a within-subjects design.

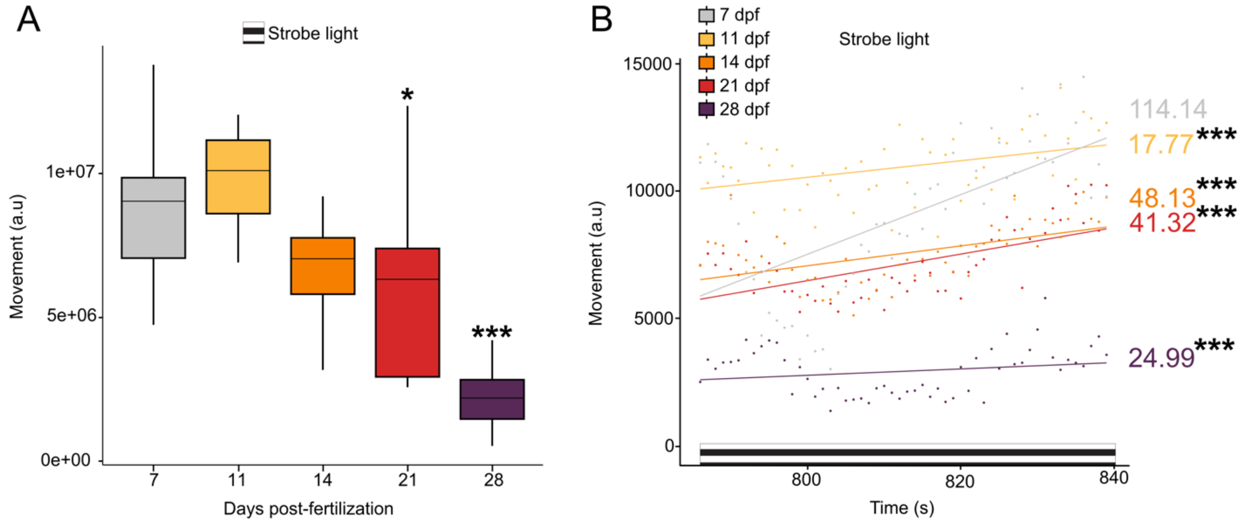


**Figure S3: Older WT-TL larvae of 28 dpf exhibit the lowest movement during strobe light stimuli and the slowest habituation. (A)** Overall total movement in strobe light phases. Older WT-TL larvae from 21 dpf exhibit less activity during strobe light stimuli than younger developmental stage larvae. *P* value computed by Tukey's honestly significant difference test on 1-way ANOVA [F(4,55) = 23.94, *P* = 1.67$^{e}$-11] **(B)** Linear regression of fish movement following strobe light stimuli. 7 dpf WT-TL larvae show rapid habituation to strobe light stimuli as measured by the slope. n = 24-36 for each time point. *p <0.05, ***p <0.001. *P* value computed by Tukey's honestly significant difference test on 1-way ANOVA [F(4,175) = 10.72, *P* = 8.55$^{e}$-8]. The correction factors used across development are as follows: 20 (7dpf), 10 (11dpf) and 5 (14dpf). These experiments were performed using a within-subjects design.

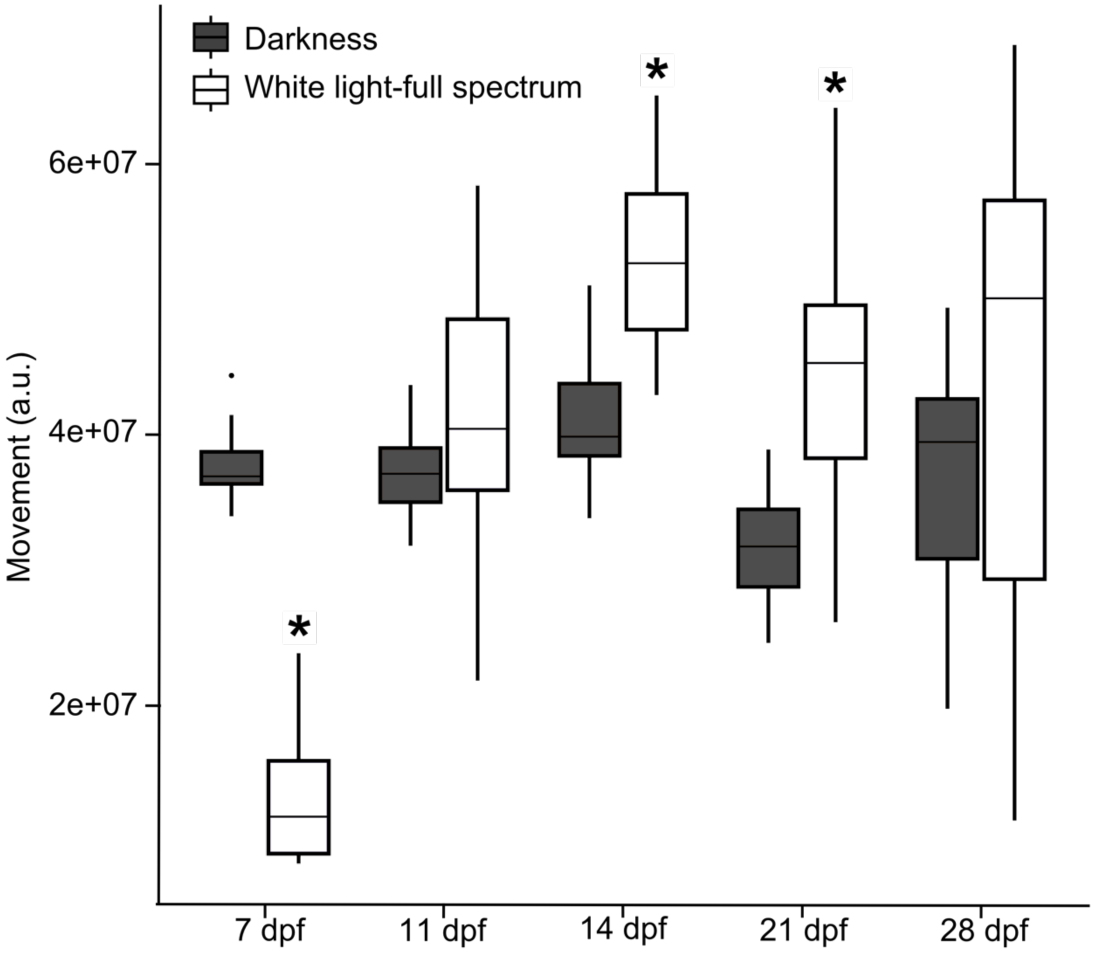


**Figure S4: Lower activity during the light phase in Light-Dark Assay in 7 dpf WT-TL larvae is reversed at 11dpf and beyond**. Total movement was quantified in both the darkness and in the 100% full spectrum white light in WT-TL animals in the Light-Dark Assay. WT-TL larvae show greater activity in the dark phases compared with light phases only at 7dpf. n = 48 – 60 animals for each time point. *p <0.05. *P* value computed by Tukey's honestly significant difference test on 2-way ANOVA [F(1,10) = 18.63, *P* = 1.02$^{e}$-11]. The correction factors used across development are as follows: 20 (7dpf), 10 (11dpf) and 5 (14dpf). These experiments were performed using a within-subjects design.

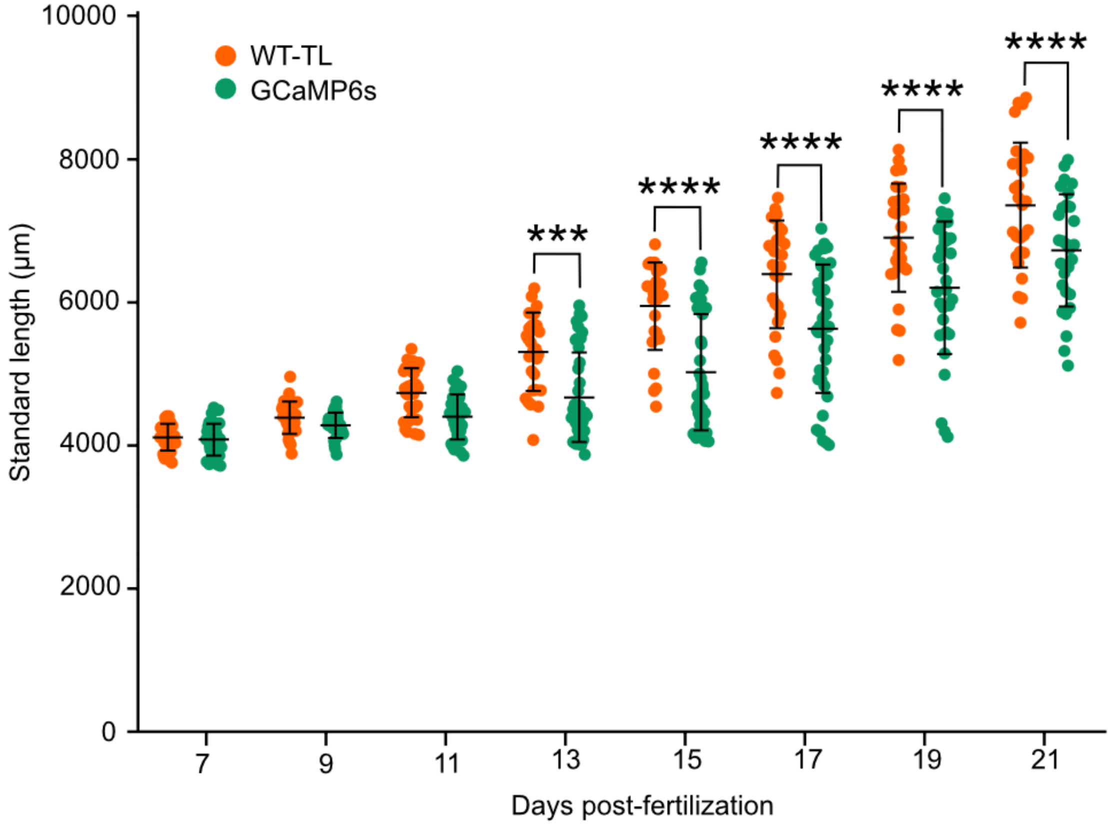


**Figure S5: GCaMP6s larvae are significantly smaller than WT-TL larvae starting from 13 dpf.** Average length of GCaMP6s and WT-TL larvae from 7 dpf to 21 dpf. Data were pooled from n=3 different batches for each line, and number of animals varies between 28-36 animals (WT-TL), and 29-41 animals (GCaMP6s). Data are shown as dots representing individual fish, and lines the mean ± SEM. *P* value computed by with Sidak's post-test on 2-way ANOVA with a significant effect of line [F(1, 564) = 148.2, *P < 0.0001*], time [F(7,564) = 200.8, *P < 0.0001*] and line and time [F(7, 564) = 7.594, P< *0.0001*]. ***p=0.001, ****p<0.0001. These experiments were performed using a between-subjects design.

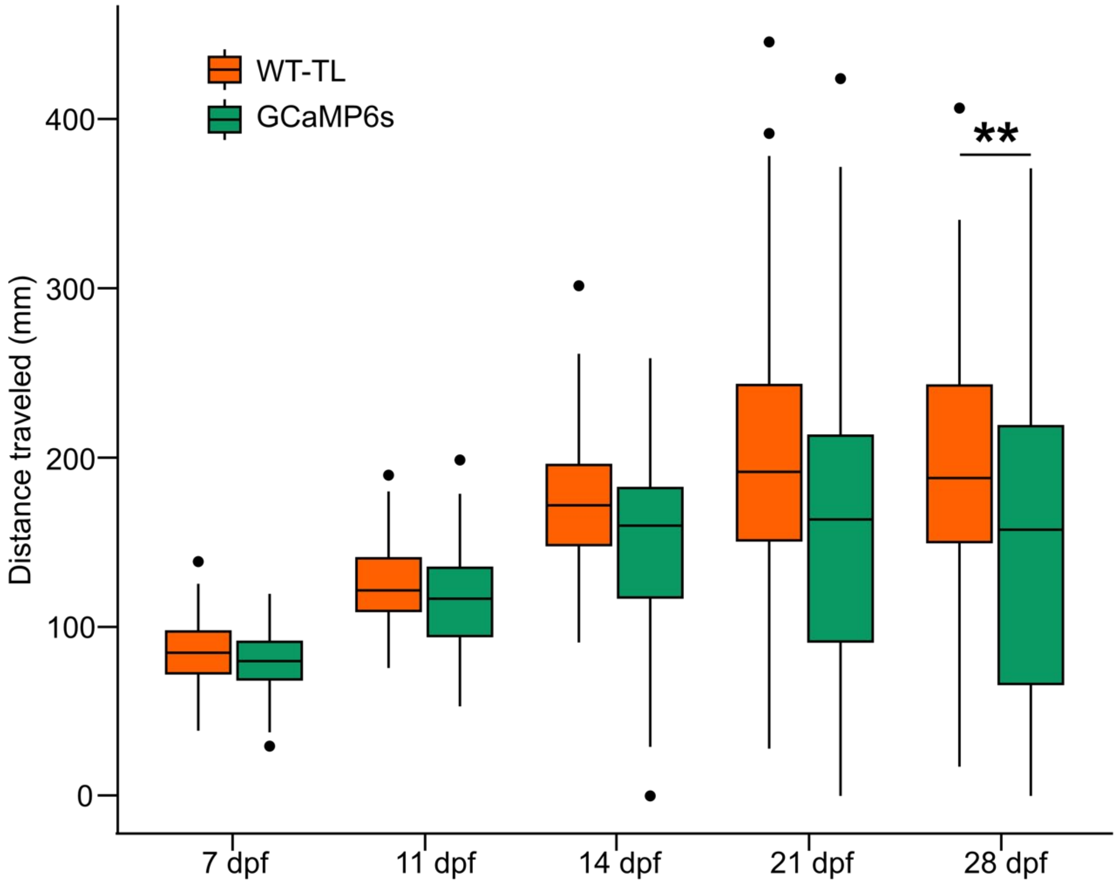


**Figure S6: Locomotion during the first 5 minutes of Light-Dark Assay at different time points.** Only 28 dpf GCaMP6s larvae show a significant decrease in the distance traveled during the first 5 minutes of Light-Dark Assay compared to the WT-TL line. N ≥ 3 per condition, n = 48 – 60 animals for each time point. *P* value computed by Tukey's honestly significant difference test on 2-way ANOVA [$F(4,590)$ = 2.234., *P =0.0519*]. **p<0.01. These experiments were performed using a between-subjects design.

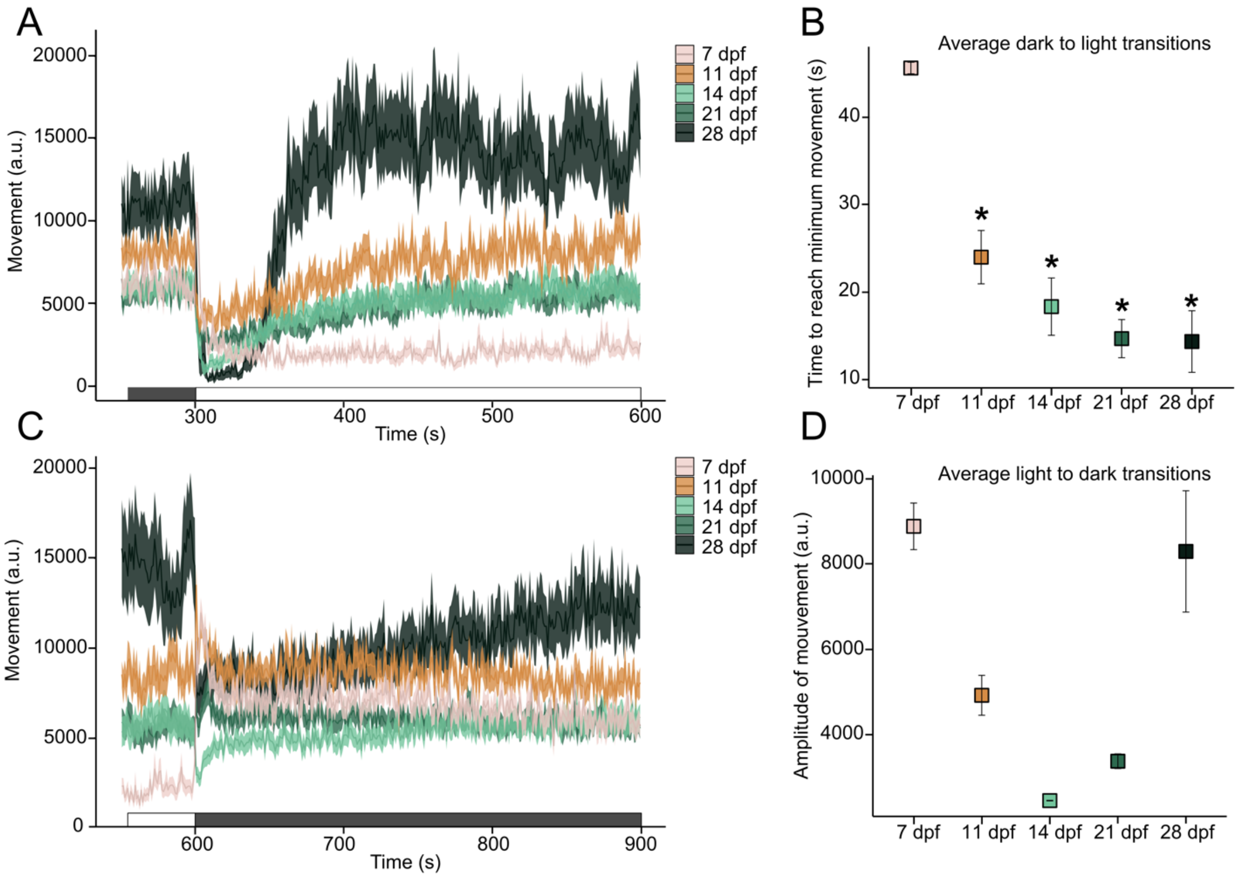


**Figure S7: Shift of sensorimotor responses to light-dark transitions in GCaMP6s line during the second week of larval development (A)-(B): Transition from dark to light:** As with the wild-type line, older GCaMP6s larvae freeze within 20 seconds of the light turning on, while 7 dpf larvae take longer to reduce their movement. Only the 28 dpf GCaMP6s fish present the sharp increase in locomotion observed in the wild-type fish following the initial freezing. *p <0.05. *P* value computed by Tukey's honestly significant difference test on 1-way ANOVA [$F(4,10) = 40.57$, $P = 3.76^{e}\text{-}7$]. **(C)-(D): Transition from light to dark.** This transition has minimal impact on older larvae, except in 28 dpf animals, where an initial burst of activity, similar to that in 7 dpf larvae, is triggered. The correction factors used across development are as follows: 20 (7 dpf), 10 (11 dpf) and 5 (14 dpf). *p <0.05. *P* value computed by Tukey's honestly significant difference test on 1-way ANOVA [$F(4,10) = 16.23$, $P = 0.0002$]. These experiments were performed using a within-subjects design.

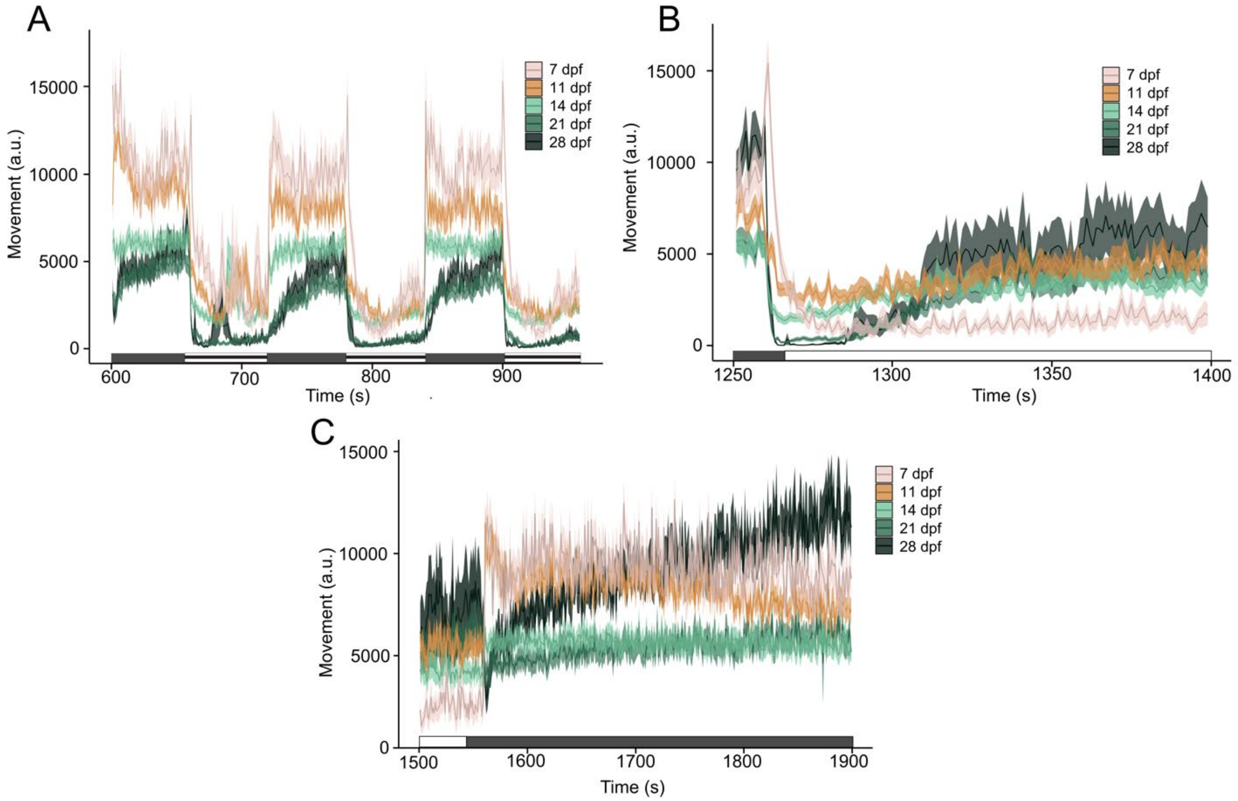


**Figure S8: Behavioral responses in the GCaMP6s line across development.** (**A**) Strobe light induces a freezing response at all stages. (**B**) At 7 dpf, a dark-to-light transition induces a burst of activity followed by a rapid decrease in activity. In older larvae, the transition leads to an immediate freezing response followed by a slow recovery to return to baseline activity. (**C**) Light-to-dark transition induces a peak activity in GCaMP6s animals until 14 dpf but leads to an abrupt reduction in movement in 21 and 28 dpf fish followed by a gradual return to baseline activity only in 28 dpf. Data are shown as the mean ± SEM, n = 24-36 animals for each time point. The correction factors used across development are as follows: 20 (7 dpf), 10 (11 dpf) and 5 (14 dpf). These experiments were performed using a within-subjects design.

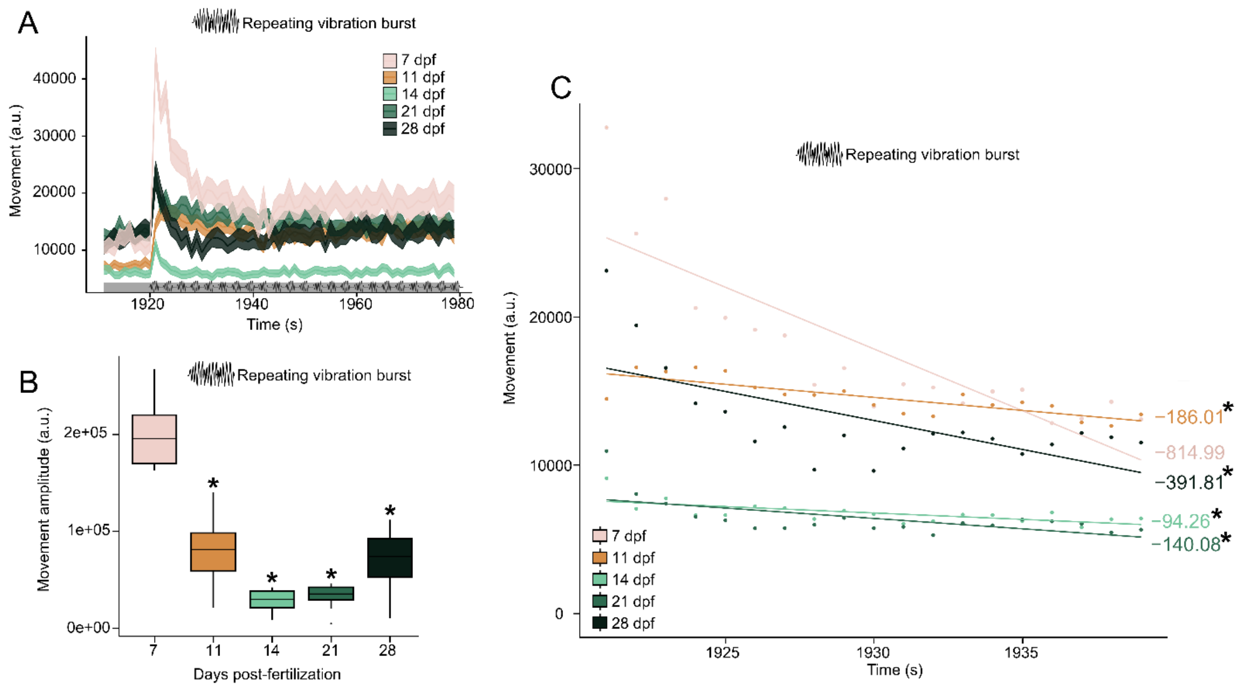


**Figure S9: Developmental U-shaped sensorimotor responses to vibratory stimuli in the Multi Stimuli Assay in GCaMP6s line.** (**A**) Repeating short bursts of vibration (400Hz) induce an initial rapid response followed by a gradual habituation to the stimuli, as observed in WT-TL larvae. (**B**) Amplitude of movement reached after the first vibration burst (400 Hz) of the habituation sequence. The amplitude was calculated by comparing the response to the vibration to the average movement during the 3 seconds immediately before the stimuli. GCaMP6s animals show the same U-shaped trend across development as observed in wild-type animals. *P* value computed by Tukey's HSD test on 1-way ANOVA [F(4,55) = 29.72, *P* = 3.53e-13]. (**C**) Linear regression of fish movement following repeating vibration burst, representing habituation. 7 dpf larvae show the highest rate of habituation. n = 24-36 animals for each time point. The correction factors used across development are as follows: 20 (7 dpf), 10 (11 dpf) and 5 (14 dpf). *p <0.05. *P* value computed by HSD test on 1-way ANOVA [F(4,55) = 41.58, *P* = 5.06e-16]. *p <0.05. These experiments were performed using a within-subjects design.

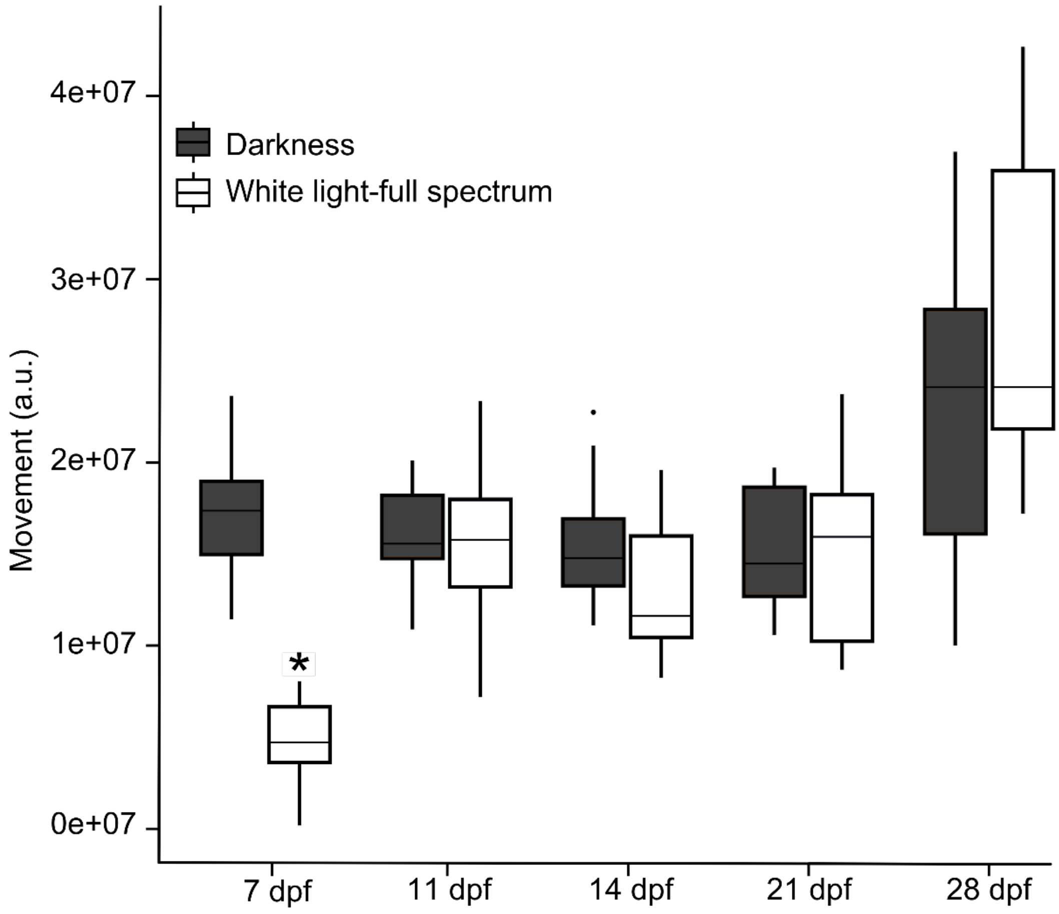


**Figure S10: Lower activity in light phase in 7 dpf GCaMP6s larvae in Light-Dark Assay while no difference is observed in older animals.** Total movement for all darkness and in the 100% full spectrum white light phases in GCaMP6s animals. As observed in WT-TL animals, GCaMP6s larvae also display greater activity in the dark phase only at 7dpf. n = 48 – 60 animals for each time point. *p <0.05. The correction factors used across development are as follows: 20 (7dpf), 10 (11dpf) and 5 (14dpf). These experiments were performed using a within-subjects design. *P* value computed by Tukey's honestly significant difference test on 2-way ANOVA with a significant effect of development stage [F(4,10) = 27.42, *P* = 8.5e-16], phase [F(1,10) = 5.7, *P* = 0.018] and developmental stage and phase [F(4,10) = 8.7, *P* = 3.05e-06].

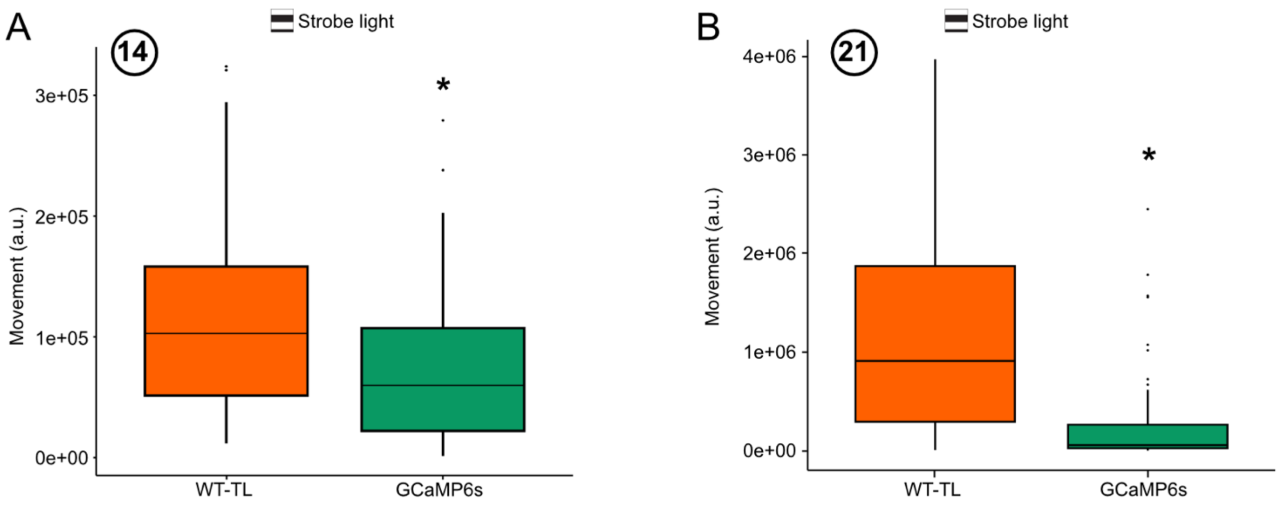


**Figure S11: Reduced activity during strobe light stimuli in GCaMP6s fish compared to WT-TL fish at 14 and 21dpf.** GCaMP6s animals exhibit a marked reduction in the total movement during the strobe-light phases compared to their wild-type counterparts at 14 dpf (**A**) and 21 dpf (**B**). n = 24 animals at 14dpf and 36 animals at 21 dpf for both lines. The correction factors used for transgenic animals is 3 for 14 and 21 dpf. *$p < 0.05$. *P* value computed by Tukey's HSD significant difference test on 1-way ANOVA for 14 dpf ; [$F(1,117) = 14.77$, *P =0.000198* ] and 21 dpf [$F(1,117) = 37.2$, *P* = 1.41e-18]. These experiments were performed using a between-subjects design.

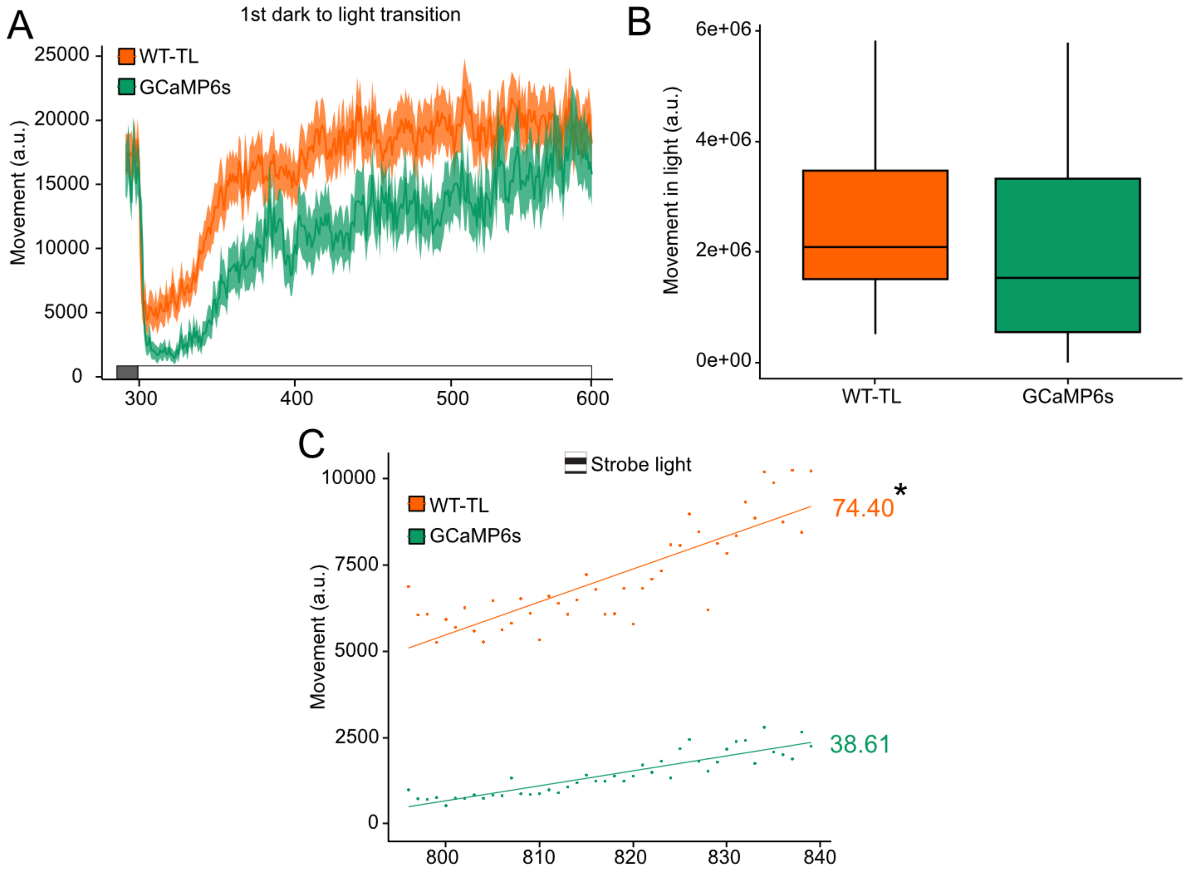


**Figure S12: Comparison of response to changes in light conditions between GCaMP6s and WT-TL larvae at 21 dpf during the Multi Stimuli Assay.** (**A**) The transition from dark to light evokes the same behavioral responses in GCaMP6s larvae and WT-TL animals. GCaMP6s show a greater initial response but a similar recovery rate. (**B**) No difference in total movement during the light phases is detected between both lines at that age. *P* value computed by Tukey's HSD significant difference test on 1-way ANOVA [$F(1,22) = 2.61$, $P = 0.118$]. (**C**) Following the initial freezing response to strobe light, animals recover as shown by an increase in movement during the later stage of the strobe phase. GCaMP6s fish show slower recovery to strobe light. *P* value computed by Tukey's HSD significant difference test on 1-way ANOVA [$F(1,355) = 6.64$, $P = 0.01$]. n = 36 animals for both lines. The correction factors used for transgenic animals at 21dpf is 3. *p <0.05. These experiments were performed using a between-subjects design.

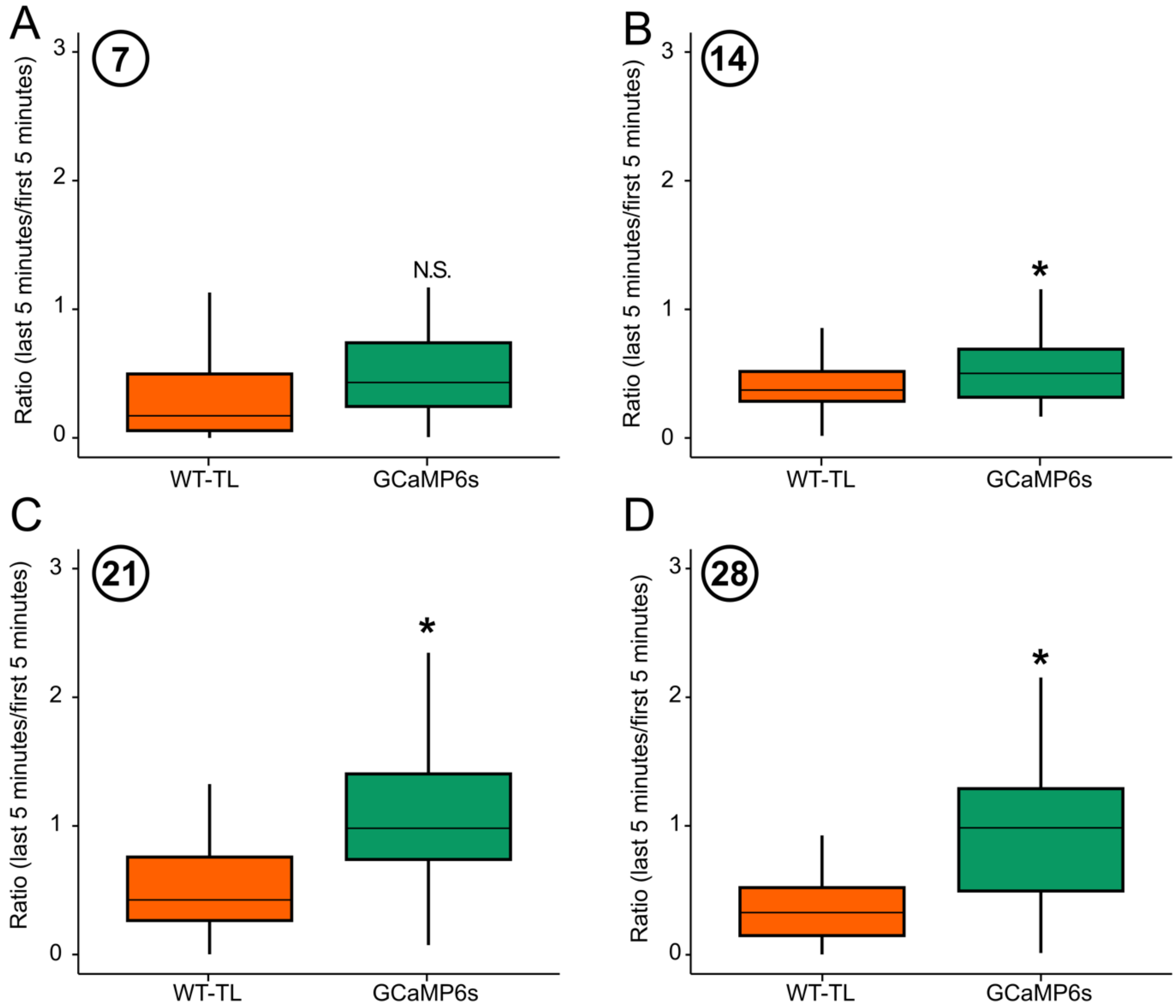


**Figure S13: GCaMP6s animals show hyperactivity at the end of the Multi-Stimuli Assay at later developmental stages.** (**A**)-(**D**) We calculated the ratio of activity between the last 5 minutes of the MSA assay compared with the first 5 minutes. A ratio greater than 1 indicates an increased activity at the end of the protocol. Overall, WT-TL larvae move less at the end of the MSA across all stages while GCaMP6s show a greater activity at 21 and 28 dpf. (**A**) 7dpf, (**B**) 14dpf, (**C**) 21dpf and (**D**) 28dpf. n = 24 – 36 animals for both lines and for each time point. The correction factors used for transgenic animals are as follows: 2 (7 and 28 dpf), and 3 (14 and 21dpf). For 7 dpf: *P* value computed by Tukey's HSD significant difference test on 1-way ANOVA [$F(1,117) = 2.407$, $P = 0.124$]. For 14 dpf: *P* value computed by Tukey's HSD significant difference test on 1-way ANOVA [$F(1,117) = 9.29$, $P = 0.002$].For 21 dpf: *P* value computed by Tukey's HSD significant difference test on 1-way ANOVA [$F(1,117) = 9.29$, $P = 0.002$]. For 28 dpf: *P* value computed by Tukey's HSD significant difference test on 1-way ANOVA [$F(1,286) = 4.84$, $P = 0.028$]**.** These experiments were performed using a between-subjects design.

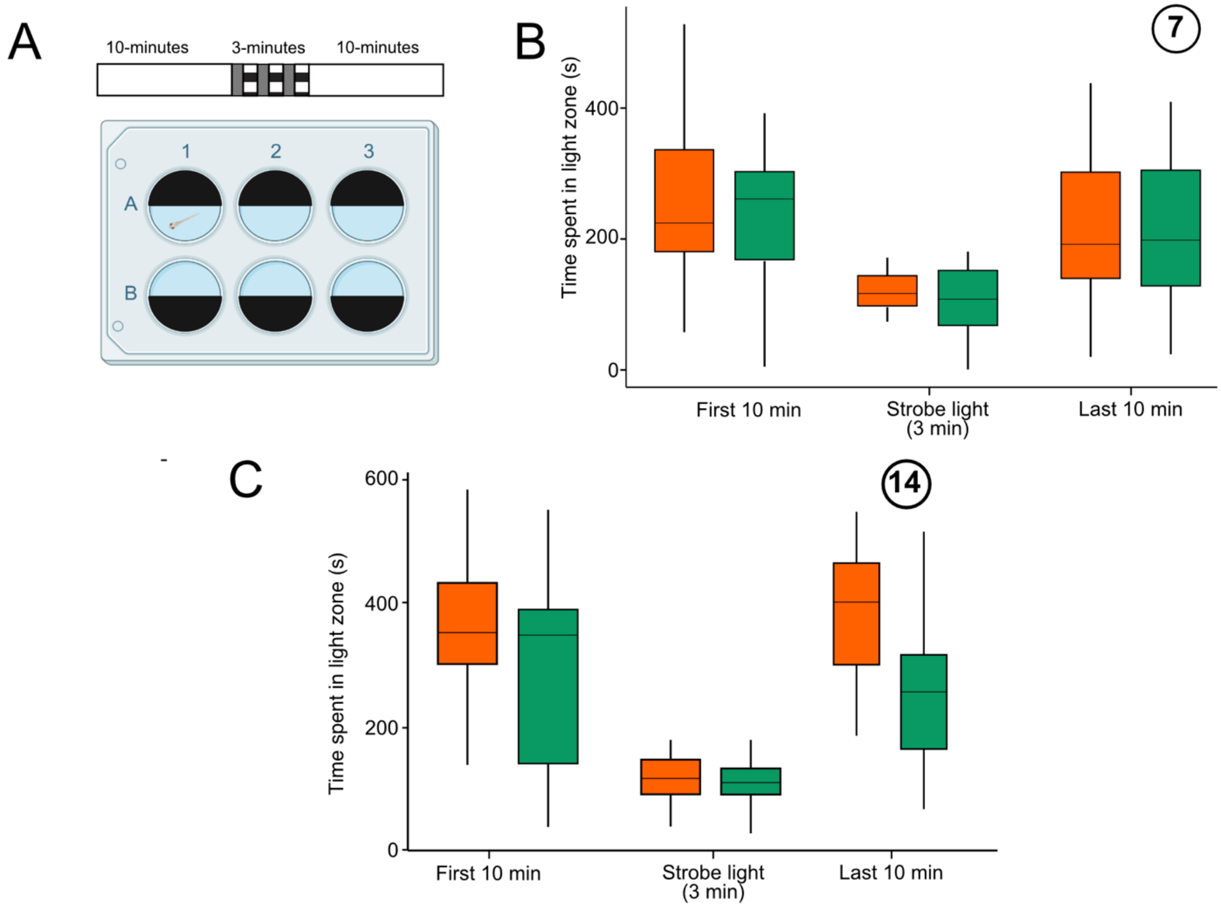


**Figure S14: GCaMP6s larvae do not show any significative difference in light-dark preference assay compared to WT-TL larvae at 7dpf.** (**A**) Overview of the assay. (**B**) Quantification of time spent in light side during the 10-minutes prior the strobe light, during the strobe light and 10-minutes after the strobe light show in 7 dpf and (**C**) 14 dpf WT-TL and GCaMP6s larvae. We did not observe a significant difference between both lines at 7dpf, but 14 dpf GCaMP6s larvae tend to move less during the last 10 minutes compared to WT-TL larvae. n = 12 animals for both lines. A correction factor of 2 was used for transgenic animals. These experiments were performed using a between-subjects design.
